# A bulk-surface moving-mesh finite element method for modelling cell migration pathways

**DOI:** 10.1101/2023.02.23.529823

**Authors:** Anotida Madzvamuse, David Hernandez–Aristizabal, Diego A. Garzon–Alvarado, Carlos A. Duque–Daza

## Abstract

Cell migration is an ubiquitous process in life that is mainly triggered by the dynamics of the actin cytoskeleton and therefore is driven by both mechanical properties and biochemical processes. It is a multistep process essential for mammalian organisms and is closely linked to development, cancer invasion and metastasis formation, wound healing, immune response, tissue differentiation and regeneration, and inflammation. Experimental, theoretical and computational studies have been key to elucidate the mechanisms underlying cell migration. On one hand, rapid advances in experimental techniques allow for detailed experimental measurements of cell migration pathways, while, on the other, computational approaches allow for the modelling, analysis and understanding of such observations. Here, we present a computational framework coupling mechanical properties with biochemical processes to model two–dimensional cell migration by considering membrane and cytosolic activities that may be triggered by external cues. Our computational approach shows that the numerical implementation of the mechanobiochemical model is able to deal with fundamental characteristics such as: (i) membrane polarisation, (ii) cytosolic polarisation, and (iii) actin-dependent protrusions. This approach can be generalised to deal with single cell migration through complex non-isotropic environments, both in 2- and 3-dimensions.

**Author summary:** When a single or group of cells follow directed movement in response to either chemical and/or mechanical cues, this process is known as cell migration. It is essential for many biological processes such as immune response, embryogenesis, gastrulation, wound repair, cancer metastasis, tumour invasion, inflammation and tissue homeostasis. However, aberrant or defects in cell migration lead to various abnormalities and life-threatening medical conditions [1–4]. Increasing our knowledge on cell migration can help abate the spread of highly malignant cancer cells, reduce the invasion of white cells in the inflammatory process, enhance the healing of wounds and reduce congenital defects in brain development that lead to mental disorders.

In this study, we present a computational framework that allows us to couple mechanical properties with biochemical signalling processes to study long time behaviour of single cell migration (either directed or random). The novelty is that the evolution law for the velocity (also known as the flow or material velocity) is described by a biomechanical force balance model posed inside the cell and this in turn is driven by the actomyosin spatiotemporal model (following the classical theory of reaction-diffusion) which is responsible for force generation as described in many experimental works [2, 5, 8, 10, 11]. Hence, our modelling approach is based on a new mathematical formalism of bulk-surface partial differential equations coupled with a novel adaptive moving-mesh finite element method to allow for significant cell deformations during migration. The approach set premises to study cell migration through complex non-isotropic environments, thereby giving biologists a predictive tool for modelling cell migration.

## Introduction

Cell motility is a phenomenon that occurs in every stage of life. It is a cyclic multi–step process in the development and maintenance of multi–cellular organisms. It consists of actin–dependent protrusions at the cell leading edge; integrin–mediated adhesions to the extracellular matrix; and acto– myosin–driven contraction of the cell [2–5]. Cells have a remarkable ability to sense both physical and chemical signals that guide them to navigate through complex environments in a process known as cell migration. This process plays a pivotal role in embryonic development [6], wound healing [7], immune response [8] and cancer metastasis [9]. Cell migration is a product of the interplay between the cytoskeleton (specially the actin cytoskeleton) and several molecules and structures such as myosin motors and focal adhesions [2,8]. The actin cytoskeleton is composed of a cross–linked array of actin filaments (F–actin), the polymeric form of actin (G–actin), and associated proteins (proteins that interact with both G–actin and F– actin) [2, 5, 8, 10, 11]. This array is mainly located at the cell cortex, beneath the cell membrane. In this region, F–actin is constantly polymerised and depolymerised.

In the last few decades research studies have explored mathematical models to help elucidate mechanisms underpinning intercellular dynamics [12]. Mechanical models can be classified by the way they treat cells. Some of them describe cell density in a continuum medium, e.g. [13, 14], where diffusive terms account for random migration, advective terms for directed migration and reactive terms for proliferation, death and other phenomena. Other models consider cells as particles that follow prescribed rules based on biological observations, such as [15]. These are known as agent- or individual-based models. Furthermore, there are models that directly study the interactions and activities of chemicals within the intra and extracellular domains as well as on the membrane [16, 17]. However, the majority of some of these studies fail to take into account important morphological (e.g. geometrical) properties of cells and their environments. Properties such as shape morphology, curvature, surface tension, among others are not quantifiable under these approaches. On the contrary, models that consider each cell as a separate geometric entity can overcome such limitations, see for example, [1, 15, 18–24, 26–29].

Our approach is a generalisation and substantial extension of this approach whereby we consider each cell as a separate geometric entity. This allows us to encode and therefore model, naturally, some of key biophysical features of single as well as collective cell migration such as (i) the biochemical interplay among different molecules in the extracellular matrix, on the membrane and in the cytosol and (ii) the mechanical response of the cytoskeleton and the membrane as a consequence of the biochemical dynamics.

Regarding the biochemical system, it needs to be able to evolve in stable and polarised states. In [30, 31] it is argued that such dynamics may be modelled as an excitable system which can switch from one steady state to another. For example, in the wave–pinning model introduced in [23] and extended in [1, 24], external stimuli switch the steady state of the system in a specific region which then triggers the evolution of a wave front that is pinned as it reaches regions where the steady state was not switched. On the other hand, Goehring et al. [32] and Meinhardt [33] suggested that such dynamics may be due to Turing instability which have the possibility to develop highly polarised states even from small perturbations. Turing instability may account for signalling amplification on the cell membrane, a key stage of cell polarisation. Following this concept, Meinhardt [33] introduced a model for cell migration able to adapt to external cues and to intensify such cues.

However, Meinhardt’s model as well as the wave–pinning do not offer a biologically real representation of the dynamics of cell polarisation since their variables are hypothetically postulated, and therefore do not directly represent specific molecular species [8]. A more relevant work is that of Cheng and Othmer [29] who introduced a biologically motivated model to simulate the polarisation signalling pathway from the activation of the receptors to the activation of RAS proteins in Dictyostelium discoideum. This model was rigorously compared with experimental results and showed great consistency. However, the implementation in a moving cell framework was not implemented as this would be too expensive in terms of computational effort.

Regarding the mechanical system, the cytoskeleton plays a major role in cell migration. The mechanical properties are highly dependent on actin filaments, microtubules and intermediate filaments. The rigidity of the cell is given by the amount and organisation of actin filaments [34]. In addition, changes on the rigidity of the extracellular matrix lead to changes in cell rigidity—a process that corresponds to cell adaptation. Cells sense external mechanical stimuli by means of focal adhesions that act as a response trigger internal signalling pathways that lead to the reorganisation of the cytoskeleton [35]. Microtubules can resist compressive forces and work as anchorage structures when attached to focal adhesions. Intermediate filaments provide structural integrity and viscoelastic properties [36]. Thus, we see that the mechanics within cells are highly dynamic since the elastic and viscoelastic properties depend on the density and organisation of the cytoskeleton.

When modelling the mechanical behaviour, it is usually assumed that inertial forces are negligible with respect to elastic and viscoelastic forces [37]. Therefore, the system is considered to be in equilibrium. Such equilibrium is defined by the shear–stress relation of the media (which could be elastic, viscous or viscoelastic), growth and surface forces. Growth forces are derived from biochemical activity and can be applied both on the surface [26–28] and in the bulk [21, 38, 39]. Surface forces comprise both external stimuli and membrane forces as a function of the mean–curvature vector.

The coupling of Meinhardt’s model to a mechanical system has been previously done in [26–28]. Neilson et al. introduced an arbitrary Lagrangian– Eulerian surface finite element method to compute biochemical dynamics and the deformation was computed by a level set method. The velocity of the surface was defined proportional to the biochemical activity. As a result, they obtained directed and random migration led by competing pseudopods. Latter, Elliott et al. applied the evolving surface finite element method [40] to solve the biochemical dynamics and coupled the mechanics by considering a force balance model posed only on the surface membrane. They considered a protrusive force proportional to the biochemical activity, a force to control cell volume, a viscous force, external forces (such as obstacles) and membrane forces (both tension and bending). Their model was also able to reproduce directed and random migration in the absence of mechanical properties inside the cell, which is a key property of our modelling approach.

On the other hand, Campbell and Bagchi [28] modelled both intra and extracellular domains utilising a fluid–dynamics approach to couple the mechanics. They used the Stokes equation and included the membrane by adding protrusive, tensile and bending forces with the immersed–boundary method. However, these models did not consider the activity of the cytoskeleton. Instead, the driving forces were applied directly to the surface and proportional to the biochemical activity on the membrane. Thus, here we present a simplistic approach to consider cytoplasm polarisation from Meinhard’s model and apply the growth forces in the bulk driven by the mechanical properties.

In this paper, we implement a computational framework that models fundamental migration properties such as: (i) random and chemotactic membrane polarisation, (ii) cytosolic polarisation, and (iii) actin–dependent protrusion. We employ the Meinhardt’s [33] reaction–diffusion model for cell orientation on the membrane and a linear diffusion–depletion model for cytosolic polarisation. In addition we model an elastic mechanical response by considering expansive and contractile forces dependent on actin concentration, curvature and area change. The proposed computational approach is based on a novel bulk–surface moving–mesh finite element method for semi–linear parabolic partial differential equations [24, 75, 76]. We note that our computational approach does not consider fitting the model to experimental data and therefore the parameters that we use are defined to demonstrate the ability of the model to exhibit experimentally observed cell migration pathways. Implementing model fitting and parameter estimation for the mechanobiochemical model as proposed is currently beyond this study due to the lack of detailed experimental datasets both for the mechanics and the biochemical processes as well as the complex nature of coupling of mechanical properties with biochemical processes on a moving domain.

Our paper is therefore organised as follows. In Section 1 we present the mechanobiochemical model for two–dimensional cell migration which is based on biological observations. Section 2 deals with the exposition of the bulk-surface moving-mesh finite element method applied to the mechanobiochemical model. The proposed numerical method is validated in Section 3. Cell migration pathways are exhibited in Section 4. In Sections 5 and 6 we discuss the implications of the bulk-surface moving-mesh finite element method applied to a mechanobiochemical model for cell migration and conclude our study by describing open problems in computational modelling of cell migration in multi-dimensions.

## 1 Formulating the mechanobiochemical model

In this section, we will present a biological description of the phenomenon as a starting-point of the mechanobiochemical model. We will describe fundamental biological features and some examples of how these occur. We will then proceed to formulate the mechanobiochemical model based on the general biophysical principles with appropriate mathematical modelling assumptions.

### 1.1 Biological motivation

A key feature of migration is the ability of cells to polarise the activity of the cytoskeleton towards a biased direction. This polarised state is characterised by higher concentration of actin filaments at the leading edge and higher concentration of myosin II at the rear edge [5]. The high concentration of actin filaments pushes the membrane, generating protrusions, while the high concentration of myosin II pulls the cytoskeleton, producing contractility. Moreover, new focal adhesions are constantly nucleated at the leading edge, while at the same time, the ones at the rear are either broken or disassembled [2, 11].

Although this might sound simple, the machinery for cell polarisation and migration is highly complex. It comprises biochemistry, mechanics and their cross-communication. For instance, in the case of directed cell migration of *Dictyostelium*, cell surface receptors coupled to G-proteins (GPCRs) trigger different signalling cascades that lead to the formation of protrusions [41], after being activated by extracellular binding molecules. For example, activated GPCRs switch on some membrane proteins of the Ras superfamily and Phosphoinositide 3–kinase (PI3K). After that, other proteins promote the accumulation of Phosphatidylinositol (3,4,5)-trisiphosphate (PIP3) at the leading edge. Then, PIP3 recruits downstream proteins that enhance the activity of the actin network at the plasma–membrane cortex of the leading edge [42]. As a result, this activity may push the membrane (a mechanical response) extending filopodia, lamellipodia or invadopodia [8]. On the other hand, the mechanical properties of the environment can be transduced into biochemical stimuli affecting the intracellular molecular interplay—a process called mechanotransduction [2]. In addition, there is no unique signalling pathway controlling cell migration. Instead, cells have several redundant and antagonist signalling pathways [11, 43, 44].

Given such complexity, modelling this phenomenon requires a large amount of data (regarding the interactions inside and outside the cell) or an understanding of the possible underlying mechanisms. In [45] the authors indicated the following approaches: bottom–up and top–down. In the former, a set of proteins and their interactions are defined based on experimental observations. Then, a mathematical model is established where each variable directly represents the considered molecules and each term describes specific biochemical reactions. Finally, the consistency of the model is tested by comparing its evolution with the experimental observations and further mathematical analysis (e.g. stability) might elucidate unknown features of the underlying mechanism. In the latter, a general mechanism (e.g. Turing instability) able to respond in a similar manner as the experimental observations is assumed. Then, mathematical analysis may indicate what sort of molecules should be present to validate the assumptions of the selected mechanism. In this work, we will take a top–down approach and thus none of the variables will relate to a specific protein. However, we will aim to mimic the evolution of the biological system.

Let us start with the membrane dynamics. As mentioned in [45, 46], the Meinhardt’s [33] model for cell orientation may be a suitable model to generate polarised states. It is one of the so called local excitation and global inhibition (LEGI) models. A key feature of these models is the difference between the diffusion rates of activators and inhibitors—that of the former are much lower than that of the latter. Within the cell, it is common that active membrane proteins diffuse more slowly than their inactive counterparts which are generally cytosolic molecules [46]. This feature is what accounts for local excitation and global inhibition. The Meinhardt’s [33] model has the following characteristics: (i) a space– dependent reaction rate that accounts for external cues; (ii) an autocatalytic activator that amplifies the deviations from the mean; (iii) a global inhibitor that restricts the amplified activator to a few regions; and (iv) a local inhibitor that destabilises steady patterns and allows for reorientation of the cell leading edge. Thus, we will consider: (i) a set of proteins that transduce external cues into internal ones; (ii) a set of proteins that activate supportive and antagonist pathways allowing them to amplify the external cues and accumulate at the leading edge; (iii) a set of homogeneously distributed proteins that prevent the amplification of the external cues at the rear; and (iv) a set of proteins mainly located at the leading edge able to breakdown the amplification which permits the system to reorient if necessary.

The polarisation of the membrane then leads to the polarisation of the cytoplasm, in particular of the actin cytoskeleton. This means that there is a flow of information between these two domains. The first assumption of our model here will be to consider that this information goes only from the plasma–membrane to the actin cytoskeleton—we assume that while the dynamics on the membrane affects the one in the cytoplasm, the former is independent of the latter. A second assumption will be that the concentration of the actin network near the plasma–membrane is proportional to the one on the membrane. This is a hypothetical assumption to be verified or refuted experimentally. It is also mathematically important since it allows us to use Dirichlet boundary conditions which directly indicate the concentration of the bulk variables on the surface. Lastly, we will consider a simple diffusion–depletion model for the general actin–network activity.

Finally, with the polarised plasma–membrane and cytoskeleton, we can consider the mechanical response. Here, the mechanics will be coupled to the biochemistry by means of forces driven by the concentration of the chemical species. The biochemistry will also be affected by the displacements through the material velocity. First, let us simplify the mechanics of the cytoskeleton by considering linear elastic deformation and plane stress behaviour [47]. Further, let us consider that no elastic energy is accumulated and, thus, the cell does not recover its shape after each step of deformation. This is following the work in [18] where the authors acknowledged that cell cytoskeleton and adhesive structures are rapidly reconstructed—this observation allows us to refer to the material model as elastic without energy accumulation instead of perfectly plastic. Second, let us couple the biochemical dynamics by considering a bulk concentration–dependent isotropic expansion. As actin filaments grow, they apply pressure to both the plasma membrane and focal adhesions [8]. We will then consider that this growth translates into mechanical stress along the filament by the linear form: *σ*_fil_ = *Eε*_fil_, where *ε*_fil_ is the strain or deformation along the filament induced by the activity of the actin network and *E* is the modulus of elasticity that linearly relates the stress and the strain [47]. Given that the actin network is highly cross–linked, we take it as a homogeneous and isotropic material; thus, following the plane–stress strain–stress relation [48] and considering no induced shear strain, the mechanical stress now reads: (*σ_a_*)_11_ = (*σ_a_*)_22_ = (*E*/(1 - *ν*))*ε_a_* and (*σ_a_*)_12_ = (*σ_a_*)_21_ = 0, where ***σ**_a_* and *ε_a_* are respectively the stress tensor (with components (*σ_a_*)_ij_, *i,j* ∈ {1, 2}) and the deformation induced by the actin network. As a third consideration, we will take the equation for the area control presented in [26]. It describes the evolution of a variable that monitors the growth of the area. We will then consider isotropic contraction with the stress tensor: ***σ_c_*** such that (*σ_c_*)_11_ = (*σ_c_*)_22_ = (*E*/(1 - *ν*))*ε_c_* and (*σ_c_*)_12_ = (*σ_c_*)_21_ = 0, where *ε_c_* is the deformation induced by the isotropic contraction, which depends on the evolution of the area control equation in [26]. We will also include membrane tension as a traction on the boundary proportional to the mean curvature vector, as previously done in [26, 27]. Finally, similarly to [18] we will include the effect of focal adhesions as elastic supports.

Given the mentioned assumptions, we consider a two–dimensional evolving domain 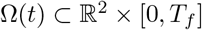 with a continuously deforming curvilinear boundary 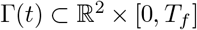 representing an evolving curve in 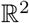. For simplicity, we ignore the nucleus and its dynamics, *Ω*(*t*) only represents the cytoplasm in the absence of the nucleus. *Γ*(*t*) is assumed to be a sharp–interface approximation of the plasma membrane. On *Γ*(*t*) we model the signal amplification using the model of [33]. In *Ω*(*t*), we model the actin–cytoskeleton dynamics by considering a linear diffusion–depletion system. Finally, we include elastic deformation as a response of the actin–cytoskeleton dynamics, the area evolution and the curvature of the membrane, respectively.

### 1.2 Formulating a surface reaction-diffusion system on the plasma membrane *Γ*(*t*)

Let us start defining the surface operators necessary to yield the reaction-diffusion equation on evolving curves. Consider an orientable curve 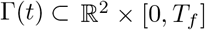 and a function 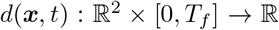 such that *d*(***x***,*t*) is the signed distance from ***x*** to *Γ*(*t*) at time *t*; then, the outward unit normal vector can be written as: ***n***(***x***,*t*) = ∇*d*(***x***,*t*)/||∇*d*(***x***,*t*)||.

Let us now define a neighbourhood of *Γ*(*t*) of size *δ* as the open subset 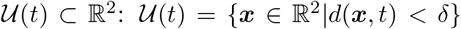. we now compute the surface gradient of a function 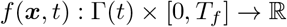, by defining an extension of this function to 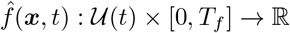 as for example:

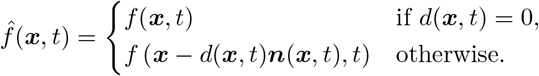

Then, the surface gradient of *f* is given by: 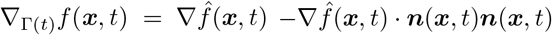. Furthermore, the Laplace–Beltrami operator is: Δ_*Γ*(*t*)_*f*(***x***,*t*) = ∇_*γ*_(*t*) · ∇_*Γ*(*t*)_*f*(***x***,*t*).

Having defined the surface gradient and the Laplace–Beltrami operator, let us continue with the derivation of the reaction-diffusion system on evolving surfaces. Let 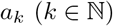 denote the *k*^th^ molecular specie resident on *Γ*(*t*) whose dynamics follow a reaction-diffusion behaviour on *Γ*(*t*), the *k*^th^ equation is given by [40, 49–51]:

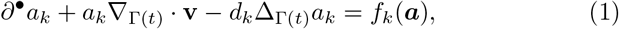

where ***a*** = (*a*_1_, … , *a_k_*) denotes the vector of the molecular species, *d_k_* is the constant diffusion coefficient of the *k*th species and **v** is the material velocity defined as: **v** = *d**u**/dt*, where ***u*** satisfies the momentum equation which will be described below Eq. (10). In addition, the material derivative is defined by: 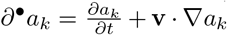. The first term of the left–hand–side of Eq. (1) is the material derivative of *a_k_*, the second term corresponds to the dilation due to the growth of the domain, and the third to the diffusion; the term of the right–hand–side represents reaction and this the only term in which interactions between the k–species take place.

For the definition of the reactive terms we will use the Meinhardt’s [33] model for cell orientation. Thus, *k* = 1, 2, 3 and the system of equations then is:

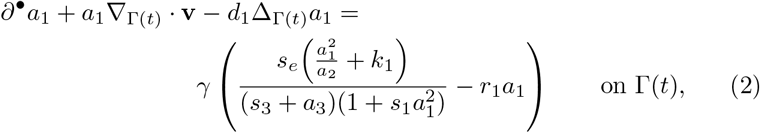

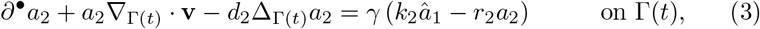

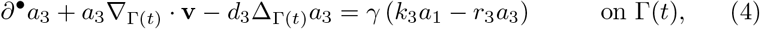

for *t* ∈ [0,*T_f_*]. With initial conditions:

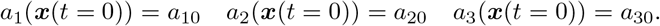

In Eqs. (2) to (4) 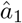 is the average value of *a*_1_; *d*_1_, *d*_2_ and *d*_3_ are diffusion coefficients; *γ* is the strength of the reaction; *k*_1_, *k*_2_ and *k*_3_ are the production rates of the activator and the global and the local inhibitor, respectively; *s*_1_ is the saturation rate of the local catalysis and *s*_3_ is the Michaelis-Menten constant; *r*_1_, *r*_2_ and *r*_3_ are respectively the consumption rate of the activator, of the global inhibitor and the local inhibitor; and *s*_e_ is the signalling parameter which captures chemotactic gradients and noise, therefore space-dependent [33]—this term can be viewed as the activity of membrane receptors.

Meinhardt [33] proposed *s*_e_ as:

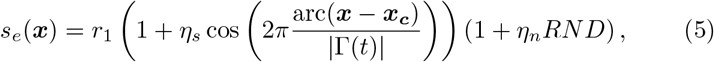

where arc(***x*** — ***x_c_***) is the distance from ***x*** to ***x_c_*** along *Γ*(*t*), ***x_c_*** is the closest point on *Γ*(*t*) to an external source indicating the directional asymmetry, *η_s_* and *η_n_* are respectively the strength of a signal coming from a specific point outside the cell and the random noise, |*Γ*(*t*)| is the size of the membrane at time *t*, and *RND* refers to a random number.

This function has two main problems: it does not model the switchable behaviour of membrane receptors and the strength of the signal does not depend on the distance to the source. As a response, Neilson et al. [26] proposed a kinetic system that models the activity of the membrane considering receptor occupancy. However, Wang and Irvine [52] indicated that receptor occupancy gradients are a consequence of the concentration gradient along the membrane of a chemoattractant. This observation, allows us to avoid the need of modelling the membrane-receptor kinetics. Instead, we modify Eq. (5) only to depend on the distance to the source. Thus, we propose the following function:

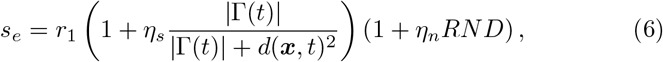

where *d*(***x***,*t*) is the distance from ***x*** to the signalling point. Section 6 illustrates the *s_e_* in the space and shows how the receptor occupancy on the membrane would behave.

**Fig 1.**
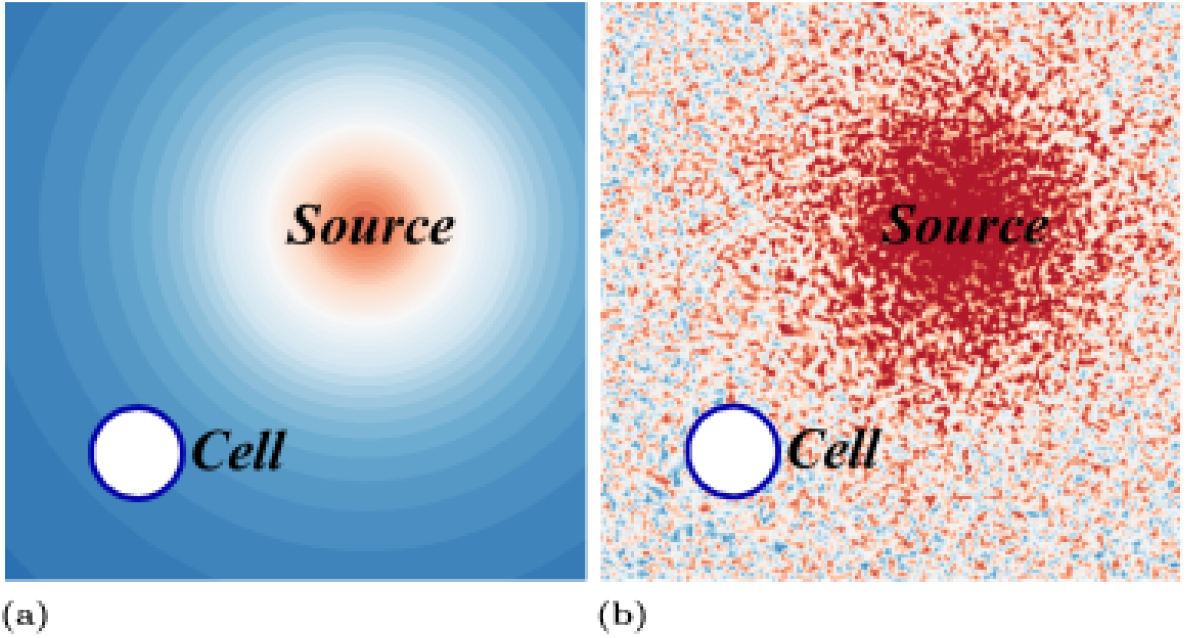
External signal *s_e_* around a circular cell with (a) *η_n_* = 0 and (b) *η_s_* = 0.

### 1.3 Formulating a bulk reaction-diffusion equation in the cytoplasm *Ω*(*t*) for actin cytoskeleton activity

The polarisation on the membrane leads to the activation of downstream signalling pathways that trigger the reorganisation of the cytoskeleton. Although we will not address directly any such signalling pathways, we will take into account the polarisation effect by assuming diffusion of a cytoplasmic chemical whose concentration at the boundary is proportional to the concentration of membrane species and a rate of degradation proportional to its concentration—again this is a hypothetical assumption. Thus, for the actin-network dynamics, let us consider a linear diffusion–depletion model with Dirichlet boundary conditions dependent on the activity on the membrane. With this boundary condition, we reproduce the polarisation of the actin cytoskeleton as a result of the polarisation of upstream regulators.

Let *a_b_* be a bulk variable representing the actin–network in the cytoplasm, then following the formulation of reaction–diffusion equations in moving domains presented in [53], the boundary value problem is given by:

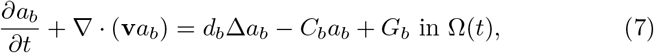

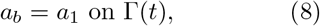

where *d_b_* and *C_b_* are respectively the diffusion coefficient and the decay rate of *a_b_* and *G_b_* is a constant production of *a_b_*.

For numerical simplicity, we will further consider that Eq. (7) reaches a steady state rapidly and thus yields:

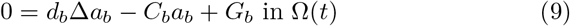

This is the final model to be solved. As can be seen, the left–hand–side of Eq. (7) is neglected.

### 1.4 Formulating the mechanical model for the cytoskeleton dynamics in the cytoplasm

As mentioned above, we will consider an elastic model with isotropic expansion and contraction due to the actin network and to the area change, respectively; adhesion to the substratum as homogeneously distributed elastic supports [18]; and surface forces due to the membrane tension [26–28]. Hence, the momentum equation [38, 39, 54, 55] reads: find the displacement field ***u***(***x***,*t*) ∈ ***C***^2^(*Ω*, [0, *T_f_*]) such that:

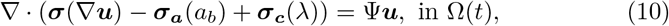

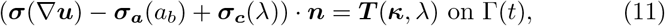

where ***σ***, ***σ_a_*** and ***σ_c_*** are respectively the true, the protrusive and the contractile stress tensors, ***Ψ*** is an elastic constant proportional to the strength of the adhesion [18], ***T*** is the tension traction on the membrane, ***n*** is the outward–pointing unit normal to the surface, λ is an area–control variable which increases if the domain grows and decreases if it shrinks [27], and ***κ*** is the mean curvature vector.

Let us now define the terms in Eqs. (10) and (11). First, the true stress tensor following Hooke’s law is given by [47]: 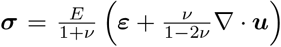, where *ε* is the strain tensor, and *E* and *ν* refer to the modulus of elasticity and the Poisson’s ratio, respectively [47]. If we adopt Voigt notation, this equation can be rewritten in matrix form, and assuming plane–stress conditions yields:

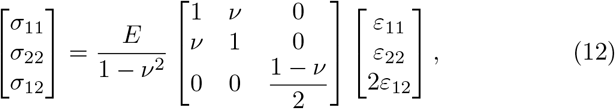

The expansion (proportional to *a_b_*) and contraction (proportional to λ) terms can be written as: ***σ**_a_* = (*E*/(1 — *υ*))_*a_b_ε*_0__***I*** and ***σ**_c_* = (*E*/(1 — *υ*))λ*ε*_0_***I***.

The area–control variable is determined by [26]:

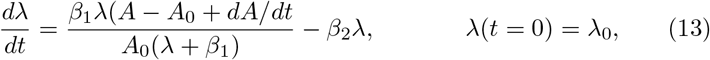

here *A*, *A*_0_ and *dA/dt* are respectively the area, the initial area, and the time derivative of the area, and *β*_1_ and *β*_2_ are positive parameters.

Finally, the mean curvature vector is [56, 57]:

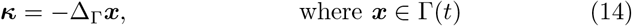

and the membrane tension is [26]:

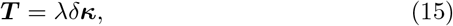

where λ is the volume constraint parameter also known as the Lagrange multiplier, *δ* is a factor of proportionality of the rigidity of the membrane.

As can be seen, Eq. (10) is a static equation. In this case, we may consider that after each computation of the biochemical model the system undergoes an elastic deformation. We may also consider that after each deformation the cell rapidly relaxes and no elastic energy is accumulated. This is equivalent to solving Eq. (10) without taking into account the prestrain given by the previous deformation step. This approach is similar to the work in [18].

Thus, our model is composed of the following differential equations: Eqs. (2) to (4), (7), (8), (10), (11), (13) and (14) that describe the evolution of *a*_1_, *a*_2_, *a*_3_, *a*_b_, ***u***, ***κ*** and λ. The full model is summarised in Table 1.

**Table 1.** Mathematical model.

| Name | Equation | Domain |
| --- | --- | --- |
| Membrane polarisation | $\partial^\bullet a_1 + a_1 \nabla_{\Gamma(t)} \cdot \mathbf{v} - d_1 \Delta_{\Gamma(t)} a_1$ | |
| | $= \gamma \left( \frac{s_e \left( \frac{a_1^2}{a_2} + k_1 \right)}{(s_3 + a_3)(1 + s_1 a_1^2)} - r_1 a_1 \right)$ | $\mathbf{x}(t) \in \Gamma(t)$ |
| | $\partial^\bullet a_2 + a_2 \nabla_{\Gamma(t)} \cdot \mathbf{v} - d_2 \Delta_{\Gamma(t)} a_2 = \gamma (k_2 \hat{a}_1 - r_2 a_2)$ | $\mathbf{x}(t) \in \Gamma(t)$ |
| | $\partial^\bullet a_3 + a_3 \nabla_{\Gamma(t)} \cdot \mathbf{v} - d_3 \Delta_{\Gamma(t)} a_3 = \gamma (k_3 a_1 - r_3 a_3)$ | $\mathbf{x}(t) \in \Gamma(t)$ |
| | $a_1 = a_{10}, a_2 = a_{20}, a_3 = a_{30}$ | $\mathbf{x}(t) \in \Gamma(0)$ |
| Kinematic model | $\mathbf{v} = \frac{d\mathbf{u}}{dt}$ | $\mathbf{x}(t) \in \Omega(t)$ |
| Cytoplasmic polarisation | $0 = d_b \Delta a_b - C_b a_b + G_b$ | $\mathbf{x}(t) \in \Omega(t)$ |
| | $a_b = a_3$ | $\mathbf{x}(t) \in \Gamma(t)$ |
| Mechanical model | $\nabla \cdot (\boldsymbol{\sigma}(\nabla \mathbf{u}) - \boldsymbol{\sigma}_a(a_b) + \boldsymbol{\sigma}_c(\lambda)) = \Psi \mathbf{u}$ | $\mathbf{x}(t) \in \Omega(t)$ |
| | $(\boldsymbol{\sigma}(\nabla \mathbf{u}) - \boldsymbol{\sigma}_a(a_b) + \boldsymbol{\sigma}_c(\lambda)) \cdot \mathbf{n} = \lambda \delta \boldsymbol{\kappa}$ | $\mathbf{x}(t) \in \Gamma(t)$ |
| | $\boldsymbol{\kappa} = -\Delta_{\Gamma} \mathbf{x}$ | $\mathbf{x}(t) \in \Gamma(t)$ |
| | $\frac{d\lambda}{dt} = \frac{\beta_1 \lambda (A - A_0 + dA/dt)}{A_0(\lambda + \beta_1)} - \beta_2 \lambda$ | $t \in [0, T_f]$ |

## 2 A bulk-surface moving-mesh finite element approach to the mechanobiochemical model in 2-space dimensions

Since we have a system of bulk–surface partial differential equations (BS-PDEs) in an evolving domain we need a robust, accurate and consistent numerical method that can compute approximate solutions of the full model. Several numerical methods have been developed and implemented to solve partial differential equations (PDEs) on evolving domains and surfaces. For example in [58] finite volumes were applied in a framework with diffusion in the bulk and on the boundary; in [59, 60] a phase–field finite–element approach was implemented to deal with a bulk–surface reaction–diffusion system; in [61] a model for the dynamics between membrane receptors and ligands was approximated by the piecewise linear coupled bulk–surface finite element method developed in [62]; in [63] the finite element approach is also implemented to solve a reaction–diffusion system able to form patterns; and in [64] by combining the virtual element method [65] and the surface finite element method [51], the authors delivered a new method to solve BS-PDEs.

In this work, we employ the bulk-surface moving mesh finite element method (BS-MFEM) to solve bulk and surface PDEs on time-dependent domains and surfaces. We assume that the mesh velocity is different from the material velocity as presented for the arbitrary Langrangian-Eulerian formulation in [25, 51, 66]. In the bulk, we will further assume that Eq. (7) is a time–independent system. Given that the coupling is through Dirichlet boundary conditions we will use the standard bulk finite element formulation. The remaining time–dependent terms will be solved using the Euler finite difference time-stepping schemes. To ensure the regularity of the finite element mesh and the convergence of the BS-MFEM on evolving domains and surfaces, we apply smoothing and re-meshing algorithms as appropriately determined by the regularity of the numerical solutions.

### 2.1 The weak formulation of the mechanobiochemical model

#### 2.1.1 The weak form of the surface reaction-diffusion system

Let us first multiply Eq. (1) by a test function *φ* and integrate over *Γ*(*t*), thus:

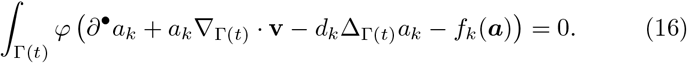

Since Green’s theorem for closed surfaces (no boundary conditions required) reads [40]:

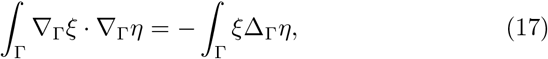

where 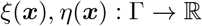, Eq. (16) can be written as:

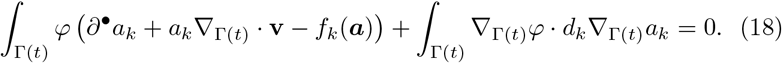

Isolating the first two terms of the right-hand side of Eq. (18) yields:

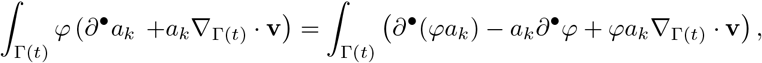

then, applying the Reynolds transport theorem [40, 51]:

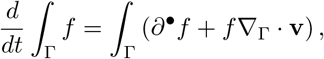

the weak form reads: find *a_k_* ∈ *H*^1^(*Γ*(*t*)) such that:

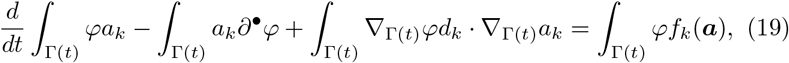

for all *φ* ∈ *H*^1^(*Γ*(*t*)).

#### 2.1.2 The weak form of the curvature equation

Let us first define the scalar product of a two second order tensors of order *n* as: 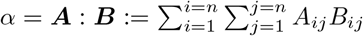. Then, multiplying Eq. (14) by a test function ***φ*** and integrating over *Γ*(*t*) yields:

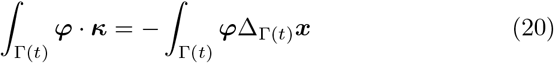

Following Eq. (17), the weak formulation reads: find *κ* ∈ *H*^1^ (*Γ*(*t*))×*H*^1^(*Γ*(*t*)) such that:

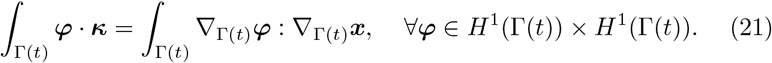

#### 2.1.3 The weak form of the bulk reaction-diffusion system

Multiplying Eq. (8) by a test function *φ*, integrating it over *Γ*(*t*) and applying Green’s theorem yields the weak problem: find *a_b_* ∈ *H*^1^(*Ω*(*t*)) such that:

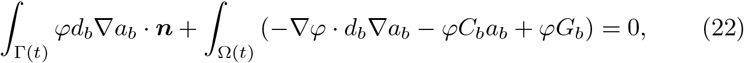

for all *φ* ∈ *H*^1^(*Ω*(*t*)).

#### 2.1.4 The weak form of the mechanical model

The weak form for Eq. (11) reads: find ***u*** ∈ *H*^1^(*Ω*(*t*)) × *H*^1^(*Ω*(*t*)) such that:

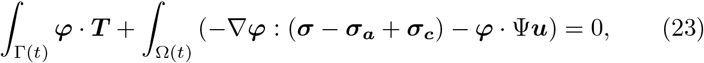

for all ***φ*** ∈ *H*^1^(*Ω*(*t*)) × *H*^1^(*Ω*(*t*)).

### 2.2 Space-discretisation

#### 2.2.1 The finite element approximation of the surface reaction-diffusion system

Let us first approximate *Γ*(*t*) by a polygonal curve 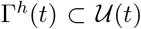 with a set of vertices 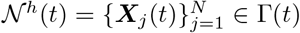 and a set of segments 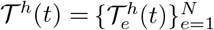. (since the curve is closed, the number of vertices is equal to the number of segments). we now define a finite element space for each *Γ^h^*(*t*) as [40]: 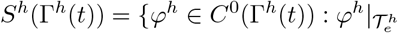 is linear affine for each 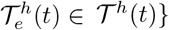, with basis functions 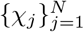.
we now write the finite element problem as: find 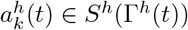 such that

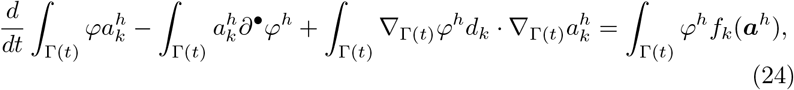

for all *φ^**h**^* ∈ *S^**h**^*(***Γ**^h^*(*t*)).

As shown in [51, 66], the discrete basis functions have the following transport property when the mesh velocity is different from the material velocity:

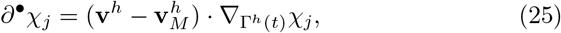

where **v**^*h*^ is the interpolated material velocity on 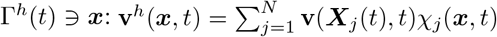, and 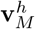 is the mesh velocity: 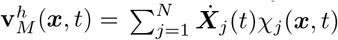. In the case that the mesh and material velocities are identical, the discrete material velocity vanishes. Since 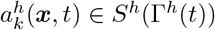 is given by: 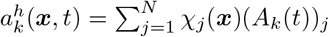, where 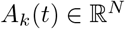 is the vector of approximated nodal values of 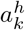 at time *t* and (*A_k_*(*t*))_*j*_ are the nodal values. Applying the transport property Eq. (25), we have the following system of ordinary differential equations:

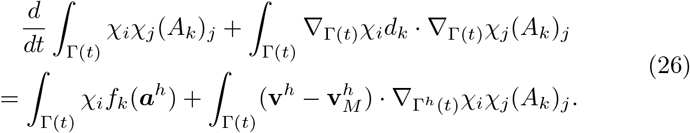

This extended ESFEM where the mesh velocity differs from the material velocity has been referred to Arbitrary Lagrangian-Eulerian evolving surface finite element method (ALE-ESFEM) [25, 51, 66].

#### 2.2.2 The finite element approximation of the curvature equation

we use the same set of vertices 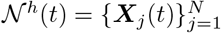 as outlined in the previous section. However, the regularity of the solutions requires us to employ at least a quadratic finite element space [56]. Therefore, we will approximate *Γ*(*t*) by a piecewise quadratic curve *Γ^q^*(*t*) (here we used the superscripts (·)^*q*^ to differentiate this approximation from *Γ^h^*) described by 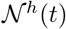 and a set of quadratic segments 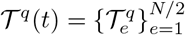. This finite element space then is: 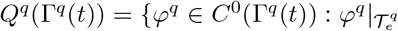 is quadratic for each 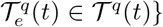. Then, the finite element problem reads: find ***κ**^q^* ∈ *Q^q^*(*Γ^q^*(*t*)) × *Q^q^*(*Γ^q^*(*t*)) such that:

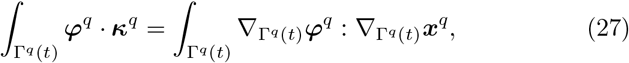

∀*φ^q^* ∈ *Q^q^*(*Γ^q^*(*t*)) × *Q^q^*(*Γ^q^*(*t*)).

#### 2.2.3 The finite element approximation of the bulk reaction– diffusion system

Let us now generate a triangulation *Ω^h^*(*t*) of *Ω*(*t*) and its corresponding finite element space. We will consider a set of nodes 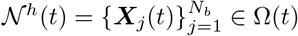 and a set of triangles 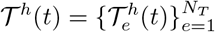 such that *∂Ω^h^*(*t*) = *Γ^h^*(*t*). The finite element space will be defined as: 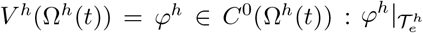 is linear affine for each 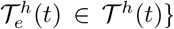. Thus, the finite element problem reads: find 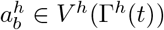 such that:

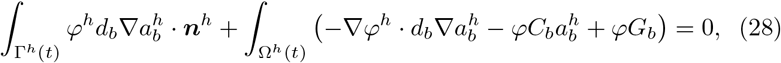

for all *φ^h^* ∈ *V^h^*(*Ω^h^*(*t*)). Since we have Dirichlet boundary conditions we set *φ^h^*|_*Γ^h^*(*t*)_ = 0, hence equation (28) now reads

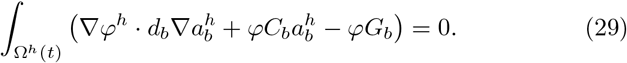

#### 2.2.4 The finite element approximation of the mechanical model

we use the same triangulation and the same finite element space as in the previous method formulate the finite element method for the mechanical model: find ***u**^h^* ∈ *V^h^*(*Ω^h^*(*t*)) × *V^h^*(*Ω^h^*(*t*)) such that:

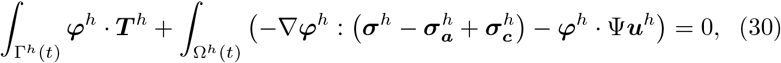

for all *φ^h^* ∈ *V^h^*(*Ω^h^*(*t*)) × *V^h^*(*Ω^h^*(*t*)).

### 2.3 Time-discretisation of the mechanobiochemical model

#### 2.3.1 The time-discretisation of the spatially discretised surface reaction-diffusion system

To solve the system of surface reaction-diffusion equations on an evolving surface, we still need to discretise the time-domain of Eq. (26). Thus, we will look for approximations of *A_k_*(*t*) that can be defined as 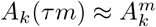, where *τ* is the time-step and 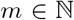. Applying a Backward-Euler time-integration scheme, we then have: find 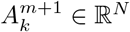 such that:

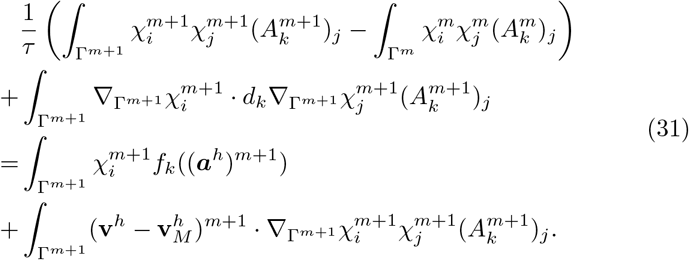

Eq. (31) in matrix-vector form is postulated as:

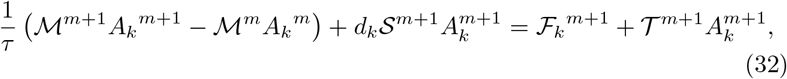

where

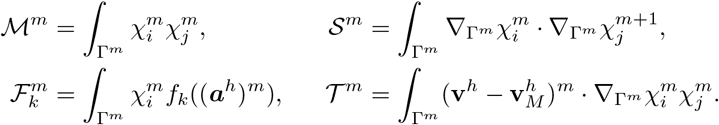

It must be noted that Eq. (3) is homogeneous along *Γ*(*t*) as long as *a*_20_(***x***) (the initial conditions) is homogeneous. In this case, the diffusion term of Eq. (31) is zero and the equation for *a*_2_ simplifies as: find 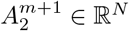 such that

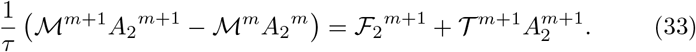

To numerically deal with the nonlinearities in Eq. (31), we employ the Newton–Raphson scheme. Since, we are using the user-defined element subroutine (UEL) of ABAQUS, we need to define the residual (Eq. (35)) and the tangent (Eq. (36)) equations. Thus, the numerical algorithm reads: find 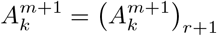 such that 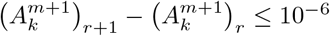 with:

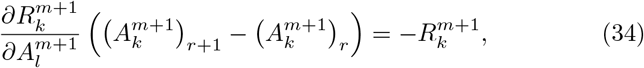

where 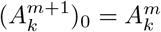. The residual equation is stated as folllows:

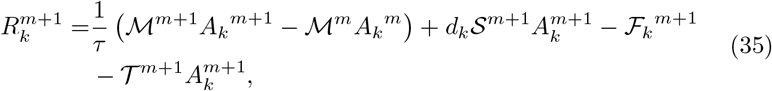

while the tangent equation states:

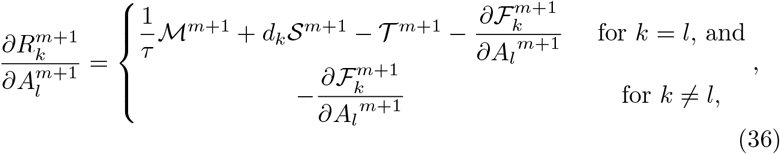

where:

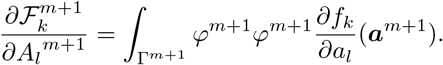

#### 2.3.2 The approximation of the area constraint *λ*(*t*)

To proceed, we now need to solve Eq. (13) for the area constraint *λ*(*t*). Here we employ the simple forward Euler scheme, thus:

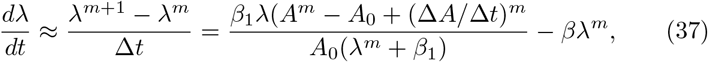

where Δ*t* can be regarded as a sensitive parameter to control the change of *λ* after each deformation. For illustrative purposes, we will set it equal to *τ*, the timestep. The terms (Δ*A*/Δ*t*)^*m*^ and *A^m^* are, respectively, the change of area and the area at time *τm*. Therefore, (Δ*A*)^*m*^ = *A^m^* — *A*^*m*-1^.

### 2.4 Mesh Smoothing and Adaptive Re-meshing Algo-rithms

Two mesh smoothing algorithms are used to improve the quality of the finite elements throughout the simulation. First, a surface mesh equidistribution scheme is implemented by means of the De Boor’s algorithm [67] and the parametric quadratic mesh. Second, the Durand’s [68] algorithm is implemented to smooth the bulk mesh. Notice that the surface mesh is made up of nodes present in the bulk mesh. Additionally, as the cell deforms continuously in space and time, a re-meshing scheme is also utilised in cases where the mesh quality drops bellow a specific threshold, monitoring, a posteriori, the mesh regularity during evolution. This re-meshing scheme is based on the Mesh2D toolbox [69, 70].

First, let us describe the mesh smoothing algorithm used on the surface. Since we defined a quadratic finite element mesh for the surface, we use the quadratic parameterisation to equidistribute the mesh. To do this, we need to define a transformation from 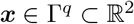 to 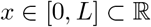, with *L* the perimeter of *Γ^q^*, such that the size of each element on *Γ^q^* is equal to the size of the image element in [0, *L*]. We might use a local transformation for *ξ* ∈ [—1,1] for each element for easy computations of the arc length. Then, we use the De Boor’s algorithm to reorganise the nodes, such that the distance between each pair of adjacent nodes is equal. After that, we use the inverse transformation to reconstruct the curve. We might use several iterations to improve the equidistribution.

Let us start by defining the local transformation and the element arc length. Let 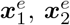 and 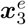 be, respectively, the start–, the end– and the middle-nodes of an element 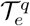. The transformation is then:

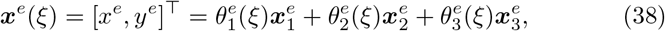

where 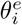 are defined as: 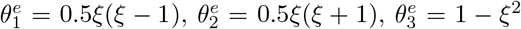. The arc length of each element 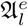 can be computed as the arc length of the parametric curve defined in Eq. (38):

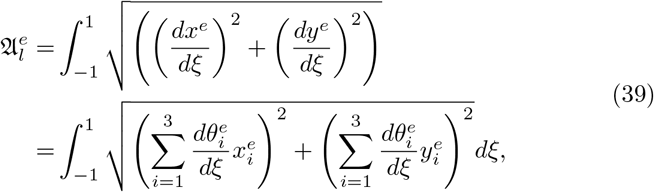

which can be approximated by Gaussian quadrature. Next, the total arc length is computed by 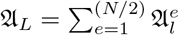. Since we want to equidistribute the surface mesh, each element must have 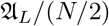 arc length and using the De Boor’s algorithm we find the appropriate node locations.

Let 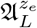 be the arc length from the first node until the end of element *e*, e.g. if 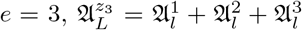. Since each element must have arc length 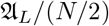, we need to find a new 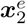, say 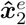, such that the new arc length is 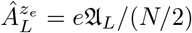. Thus, by De Boor’s algorithm we need to find and element *r* such that: 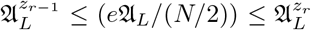. This means that 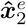 belongs to element *r*. Then, we find: 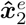 such that:

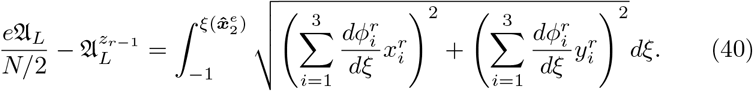

Again we use Gaussian quadrature to approximate 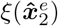, although we need to iterate several times. Once we have 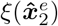, we use Eq. (38) on element *r* to find 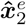.

Using this procedure, we set an equidistant mesh. However, since we are working on a closed curve, there are infinite possible meshes. For the purpose of uniqueness, we add an additional constraint such that the distance between the initial mesh and the final mesh is minimal. Let us first recall the initial mesh on [0,*L*] with the set of points 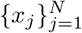 and the target set of points 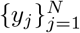. we then establish the optimisation problem as: find 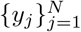 such that:

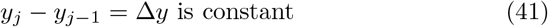

and

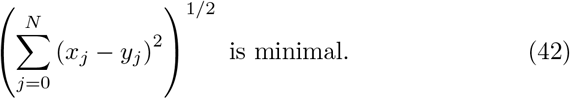

By De Boor’s algorithm we satisfy Eq. (41) and, acknowledging that defining *y*_0_ we only find one equidistant mesh, we set the target function as:

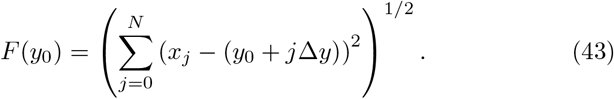

This attains its minimum value when

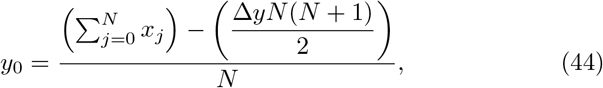

which can be negative since *Γ^q^* is a closed curve. Thus, the algorithm for equidistribution on the curve with minimal mesh displacement is as follows: (i) set a mapping from *Γ^q^* to [0, *L*], (ii) define Δ_y_ = *L/N*, (iii) establish *y*_0_ with Eq. (44), (iv) use De Boor’s algorithm to compute the other nodes, and (v) apply the inverse mapping with the Gaussian quadrature to solve Eq. (40). A schematic representation is shown in Section 6.

**Fig 2.**
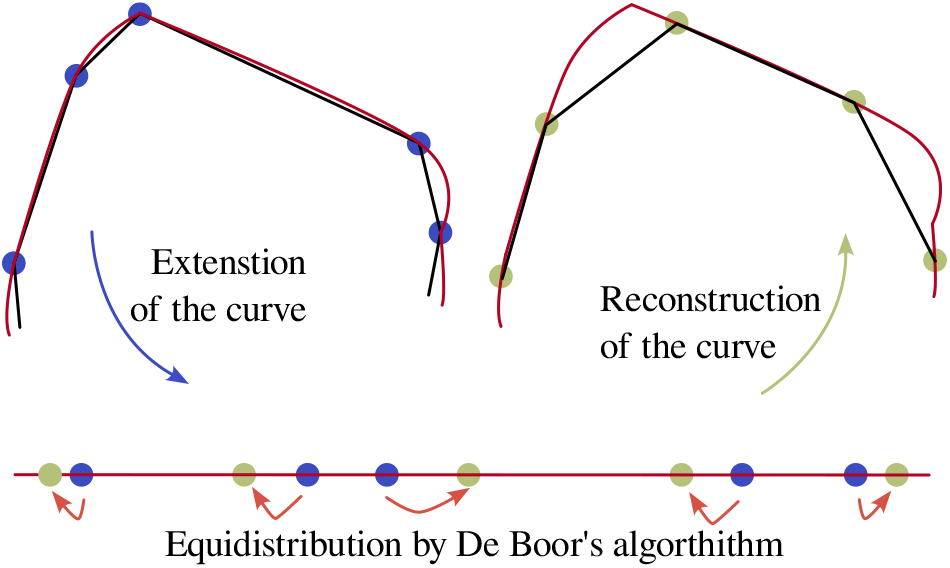
Surface smoothing scheme.

Now, for the bulk mesh smoothing we use the method introduced in [68]. We also adopt their quality metric, which for a triangle element is given by [68]:

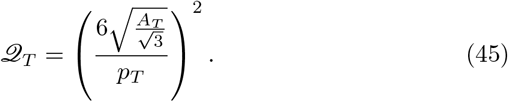

In the case that this method fails in improving the minimum quality, 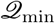, above some threshold, say 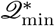, we use the Mesh2D toolbox [69, 70] to remesh the domain. Since the bulk variables are not time-dependent, it is not necessary to map the variables after the re-meshing scheme. Additionally, the equidistant surface mesh is provided as an input to Mesh2D and remains unmodified. Therefore, the surface variables do not need to be mapped either. The full algorithm can be summarised as follows:

#### The algorithm

1. Set the initial mesh

a. Generate an initial piecewise linear mesh on the surface with a even number of nodes.
b. Use the Engwirda’s [69, 70] toolbox Mesh 2D to create a piecewise linear triangular mesh.
c. Create a quadratic set of surface elements using the same nodes of Item 1a.
d. Use the Durand’s [68] algorithm to smooth the mesh.
2. Solution of field variables at time *nτ*.

a. Approximate *a*_1_ and *a*_3_ through Eq. (35).
b. Approximate *a*_2_ through Eq. (33).
c. Approximate *a_b_*, ***u*** and ***κ*** through Eqs. (27), (29) and (30).
d. Update the nodal positions as ***x**^n+1^* = ***x**^n^* + ***u**^n^*.
e. Approximate *λ* through Eq. (37).
f. If *nτ* = *T_f_* go to Item 4.
3. Improve mesh quality

a. Use the De Boor’s [67] algorithm to equidistribute the the surface mesh.
b. Use the Durand’s [68] to smooth the solid mesh until the mesh change by L_2_-norm is less than *Ite_tol_* or the maximum number of iterations *Ite*_max_ is reached. If 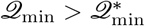 go to Item 2 if not go to Item 1.
4. Finish.

## 3 Validation of the bulk-surface finite element method

### 3.1 Numerical validation of the bulk-surface moving-mesh finite element approach

Next, we will test the convergence of the the bulk-surface moving-mesh finite element approach. For that, we run a chemotactic simulation such that the centroid of the domain moves form (0, 0) to (0, 0.5) with several time-steps and number of nodes at the boundary. We will compute the norm of the displacement as a function of time, that is: 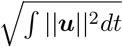, and then compute the relative error, assuming that the simulation with the smallest time step and the largest number of nodes at the boundary is the best.

We then proceed to investigate the response of the Meinhardt’s model on a stationary two-dimensional closed hypersurface. The idea here is to test the response with several combinations of parameters in terms of signal amplification and adaptation to new signals.

We exhibit results of the whole computational algorithm under two different combinations of parameters. One of them corresponding to chemotactic migration and the other to completely autonomous random movement. These numerical tests are computed using the numerical methods presented in the previous section in a desktop computer with a quad–core processor with 2.40GHz of base frequency and 32 Gb of RAM.

#### Convergence of the bulk-surface moving-mesh finite element method

Table 2 illustrates the relative errors, these reduce as the time-step becomes smaller while the number of nodes becomes larger. From these results, we will take *τ* = 0.001 and 2000 nodes to define the initial contour.

**Table 2.** Convergence of the displacement field. Relative error between the norm of the displacements for simulations with different time-step and number of nodes at the boundary.

| | $\tau = 0.002$ | $\tau = 0.0005$ | $\tau = 0.0002$ | $\tau = 0.0001$ |
| --- | --- | --- | --- | --- |
| $N = 1000$ | 8.33140e−03 | 7.76661e−03 | 7.66300e−03 | 7.64941e−03 |
| $N = 2000$ | 9.96355e−04 | 6.37406e−04 | 5.62171e−04 | 5.21905e−04 |
| $N = 4000$ | 5.22995e−04 | 1.97342e−04 | 6.98492e−05 | 0.00000e+00 |

### 3.2 The Meinhardt’s model for cell polarisation

Let us fix *a*_10_, *a*_20_, *a*_30_ *γ*, *k*_1_, *k*_2_, *s*_3_, *r*_1_, *r*_2_, *r*_3_, *η_s_* and *η_n_* (see Table 3) and vary *k*_3_, *s*_1_, and *d*_3_ (see Table 4) using the proposed parameters in [26–28, 33] for illustrative purposes. For this experiment, the domain is a circle of radius 1 centred at (0, 0), the finite element discretisation is composed of 2000 nodes and elements and the time-step is 0.001. The signal is set at (5, 5) from time 0 to 15 and at (−5, −5) from 15 to the end of the simulation. we then consider the following measures: the number of peaks *N_p_*, the highest value of the activator 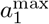, the width *w* of the peaks above 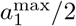 and whether the model is able to adapt to a new signal. *N_p_* allows us to determine the ability of the model to yield a polarised state, 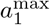 indicates the strength of the polarisation and w defines how localised it is.

**Table 3.**
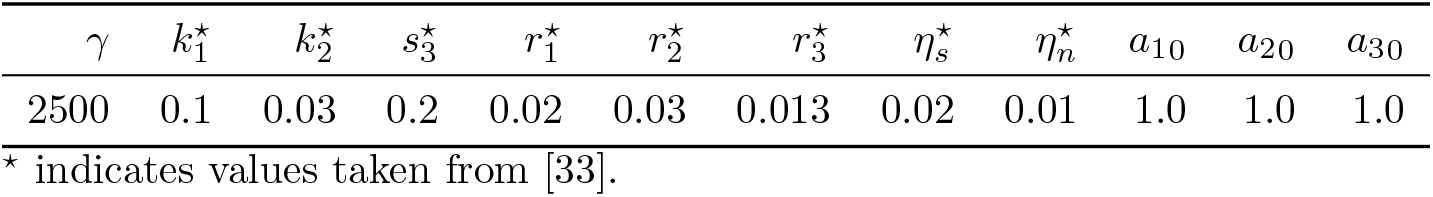
Fixed parameters for the Meinhardt’s model for cell polarisation.

**Table 4.** Meinhardt’s model response. Summary of the response of the Meinhardt’s model for cell polarisation with different parameters.

| Experiment | $k_3$ | $s_1$ | $d_1$ | $d_3$ | $N_p$ | $w$ | $a_1^{\max}$ | Adaptation |
| --- | --- | --- | --- | --- | --- | --- | --- | --- |
| 1 | 0.005 | 0.005 | 10 | 60 | 1 | 2.218 | 6.91 | Yes |
| 2 | 0.005 | 0.005 | 10 | 30 | 1 | 2.441 | 5.69 | Yes |
| 3 | 0.005 | 0.005 | 10 | 20 | 1 | 2.677 | 4.94 | Yes |
| 4 | 0.005 | 0.005 | 10 | 100 | 1 | 2.127 | 7.72 | Yes |
| 5 | 0.005 | 0.005 | 10 | 50 | 1 | 2.259 | 6.6 | Yes |
| 6 | 0.005 | 0.005 | 10 | 15 | 1 | 2.934 | 4.4 | Yes |
| 7 | 0.0025 | 0.005 | 10 | 30 | 1 | 2.975 | 8.1 | Yes |
| 8 | 0.0025 | 0.005 | 10 | 20 | 1 | 3.299 | 7.18 | Yes |
| 9 | 0.0025 | 0.005 | 10 | 15 | 1 | 3.654 | 6.49 | Yes |
| 10 | 0.005 | 0.0025 | 10 | 30 | 1 | 2.262 | 6.48 | Yes |
| 11 | 0.005 | 0.0025 | 10 | 20 | 1 | 2.466 | 5.58 | Yes |
| 12 | 0.005 | 0.0025 | 10 | 15 | 1 | 2.680 | 4.96 | Yes |
| 13 | 0.005 | 0.01 | 10 | 30 | 1 | 2.717 | 4.78 | Yes |
| 14 | 0.005 | 0.01 | 10 | 20 | 1 | 3.135 | 4.15 | Yes |
| 15 | 0.005 | 0.01 | 10 | 15 | 1 | 3.415 | 3.68 | Yes |
| 16 | 0.005 | 0.005 | 1 | 2.5 | 2 | 1.949 | 6.46 | No |
| 17 | 0.005 | 0.005 | 1 | 2 | 2 | 2.079 | 5.94 | No |

### 3.3 Chemotactic-driven directed cell migration

For the solution of the complete model, let us set the parameters of the mechanical model as presented in Table 5. These parameters were in order to validate the general ability of the computational framework to reproduce key features and properties observed during single cell migration and are not experimentally determined. A more rigorous selection would require: (i) the selection of a specific type of cell, (ii) the substratum attachment strength to define Φ, (iii) its elastic properties *ν* and *E*, the calibration of *β*_1_ and *β*_2_ to determine the velocity of the response of the area control mechanism, (iv) the membrane tension to properly define *δ*, and (v) the relation between the formation of actin filaments and the mechanical stress induced to set *ε*_0_. We defer such an experiment for future studies where we will also try and recover some of these parameters through inverse approaches for parameter estimation, using Bayesian methods [71].

**Table 5.** Parameters of the mechanical model. The last three parameters, namely 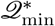, *Ite*_max_ and *Ite*_tol_, are parameters of the mesh smoothing algorithm presented.

| $\Psi$ | $\nu$ | $E$ | $\delta$ | $\beta_1$ | $\beta_2$ | $\lambda_0$ | $\varepsilon_0$ | $\mathcal{Q}_{\min}^*$ | $Ite_{\max}$ | $Ite_{\text{tol}}$ |
| --- | --- | --- | --- | --- | --- | --- | --- | --- | --- | --- |
| 0.1 | 0.45 | 1 | 0.001 | 0.2 | 0.001 | 2.0 | 2.5e−5 | 0.55 | 10 | 0.001 |

For chemotactic cell migration, we choose the parameters for Experiment 8 (in Table 4) as explained in the next section. The initial domain is bounded by a circle of radius 1 centred at (0, 0). Furthermore, the system is tested under an initial signalling point at (5, 5) that is then changed at *t* = 75 to (0,10). For this simulation, the time-step *τ* = 0.001, the surface was discretised with 2000 equidistant nodes and the size of the bulk mesh was constraint to 0.25.

#### Spontaneous random cell migration

In this case, the mechanical model is implemented with the same parameters as presented in Table 5. The initial domain is also bounded by a circle of radius 1 centred at (0, 0) and for the Meinhardt model we use the parameters corresponding to the Experiment 17—again, this choice is explained in the next section. We expect to see unsteady and completely spontaneous random cell migration using these parameters and letting *η_s_* =0 and *η_n_* = 0. The initial mesh and the time-step are the same as in the chemotactic-driven directed cell migration experiment.

## 4 Exhibiting single cell migration pathways using the bulk-surface moving-mesh finite element method

### 4.1 The Meinhardt’s model for cell polarisation

Table 4 shows the results of our experiments. We see that Experiments 16 and 17, where *d*1 = 1 and *d*2 = 2, 2.5, had a very different response compared to the others. The system developed two coexisting peaks and was not able to adapt to changes in the signal, whereas in the other experiments the system developed one peak and was able to reorient. In experiments 1 to 15, *d*_1_ = 1 and *d*_3_ varied from 15 to 100. Increasing *d*_3_ made 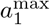 large, however, *w* decreased and the system was slower to move from one polarised state to another. A reduction in *k*_3_ from 0.005 to 0.0025 provided high values of 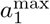 and *w*, and decreased the time from one polarised state to another; but, a further reduction, to for example 0.00125, almost homogenised the response—in other words, 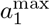 rapidly decreased. Finally, in our experiments, decreasing the value of *s*_1_, increased 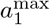 and decreased *w*.

From these results, the parameter values of Experiment 8 will be used for chemotactic-driven directed cell migration. In this case, the system is able to adapt to a new signalling source and the strength and width of the peak (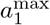 and *w*) were among the largest. This shows the ability of the system to amplify the external signal. Section 6 shows the results of the simulation with such parameters. As can be seen, the system took around *t* = 1.1 to reach a stable polarised state. After that, at time 15 the signal was changed to the opposite direction and the system needed about *t* = 150 to developed the new polarised state. The transition from *t* = 1.1 to *t* = 150 occurred as a travelling wave as indicated by the results at time 90.

### 4.2 Chemotactic-driven directed cell migration

Next, we analyse the impact of chemotactic cell migration as shown in Section 6. The simulation ran until *t* = 205 and the cell migrated towards the signalling-source points. we see the ability of the model to mimic migration pathways to specific locations and to adapt to the repositioning of the signalling sources. Section 6(a) shows the changes in shape of the cell and the position of its centroid as well as the *a_b_* field. In this case, the domain conserved a round-like shape throughout the whole simulation. It must be noticed that the cell membrane—as well as the cytosol—rapidly polarised towards the first signalling-source point (5, 5). After the change

**Fig 3.**
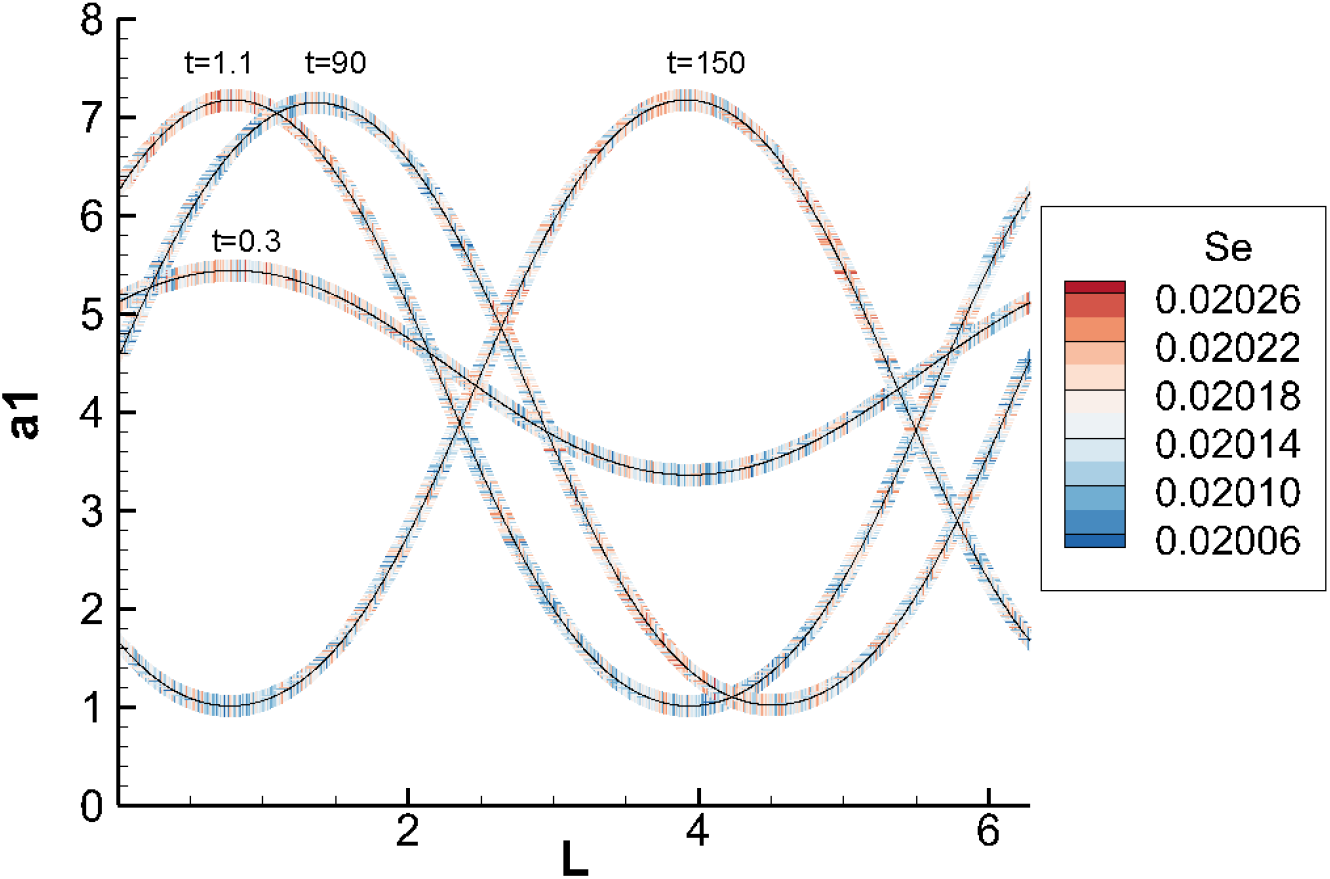
Response of the Meinhardt model. A stationary circle of radius 1 centred at (0, 0). The graph shows *a*_1_ along the arc length at simulation times 0.3, 1.1, 90 and 150. The contour plot refers to *s_e_* which started with a chemotactic signal at (5, 5) and changed to (−5, −5) at time 15. As can be seen, the system went from a polarised state around *L* = 1 to another polarised state around *L* = 4 due to the change of the signalling-source point.

in position of the signalling–source point, the cell polarised towards (10, 0) as expected. We also see that, after reaching the second point, the cell started to move around the source point. In addition, the simulation shows the appearance of a small protrusion at the rear, see the distribution of the signed curvature in Section 6(a). Section 6(b) shows the changes in area of the cell during the simulation. As can be seen, it varied strongly at the beginning, before *t* = 20, and then began to oscillate between time 3.15 and 3.22.

Finally, Section 6 shows the mesh quality (see Eq. (45)) and the positive impact of the mesh smoothing and re-meshing schemes. Section 6(a) shows, from top to bottom, the maximum, the average and the minimum element qualities, respectively, throughout the simulation. The maximum was 1 at every step which is in fact the maximum possible value. The average remained around 0.93—this is a high quality value and indicates that the mesh throughout the simulation was almost optimal. In contrast, the minimum quality fluctuated somewhat slightly with the lowest value at about 0.58 and the highest at about 0.68. In terms of the smoothing scheme, in those cases where the algorithm was sufficient to deliver an acceptable mesh, 3 or 4 iterations were needed. Furthermore, it was not necessary to use the re-meshing scheme since the mesh was at optimal regularity (for example, for 18,000 timesteps). Hence, the re-meshing algorithm was employed it was absolutely necessary, thereby saving significantly on the computational costs and efficiency of the algorithm.

**Fig 4.**
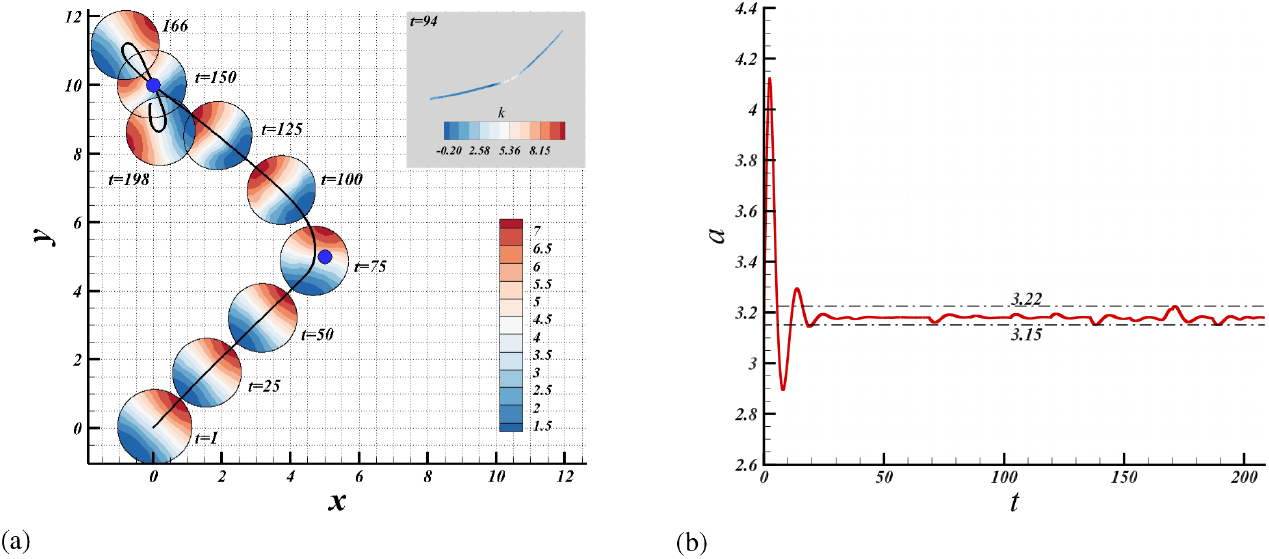
Results of chemotactic cell migration. (a) Results of the chemotactic-driven cell migration at *t* = 1, 25, 50, 75, 100, 125, 150, 166, 198 and the migration locus of the centroid. The colour map indicates the *a_b_* field, the blue dots are the signalling-source points and the continuous and solid black line is the trajectory made by the cell centroid. The inner box indicates the curvature of the rear of the cell membrane at *t* = 94. The colour map refers to the signed curvature. (b) Illustration of changes in area of the cell during for chemotactic-driven cell migration.

**Fig 5.**
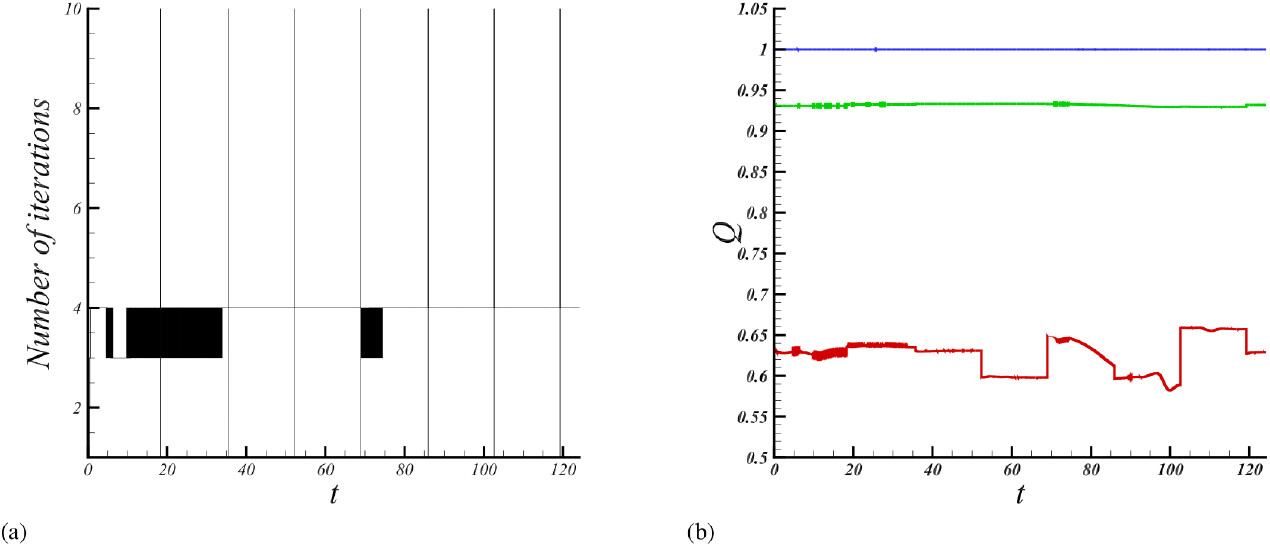
Report of mesh statistics and smoothing iterations at each step during chemotactic-driven cell migration. (a) Mesh quality statistics. From top to bottom the curves indicate respectively the maximum, the average and the minimum qualities. (b) Number of smoothing iterations. The vertical lines indicate re-meshing steps.

### 4.3 Spontaneous random cell movement

Section 6 and Section 6 show the results for the case of random cell migration. In this case, the simulation was run until *t* = 124 and the cell migrated without any external signal or space dependent parameter. Section 6(a) shows the trajectory of the cell centroid that indicates zigzag migration and some directional persistence—rather than moving around (0, 0) the cell moved towards (0.6, —1.6). In Section 6, the shape and the *a_b_* field is illustrated at *t* = 1, 25, 50, 75, 100, 124. Although the deformation of the domain is higher than in the previous case, it is noticeable how the round-like shape is still dominant. In addition, the commonly reported behaviour of the Meinhardt model can be seen, i.e., the appearance of two competing pseudopods [26–28, 33]—nonetheless, in this simulation there is no external stimuli at all. Further, Section 6(b) shows that the area of the 724 cell fluctuated much more, between 2.7 and 3.53 after *t* = 20.

**Fig 6.**
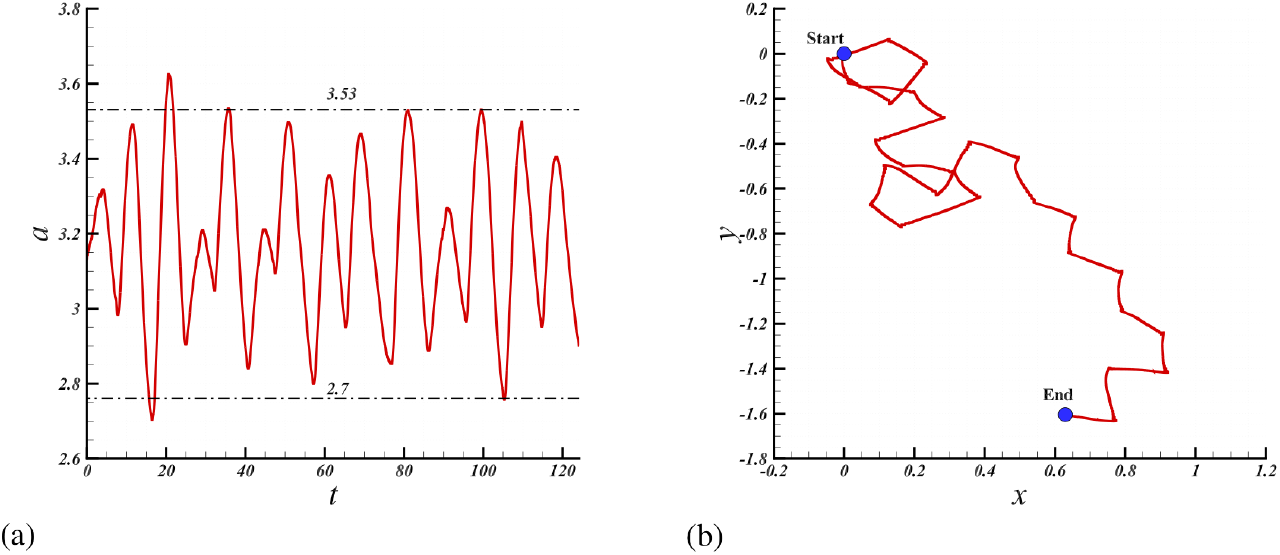
Results of spontaneous cell migration. (a) Tracking of the centroid of the cell and (b) area evolution.

**Fig 7.**
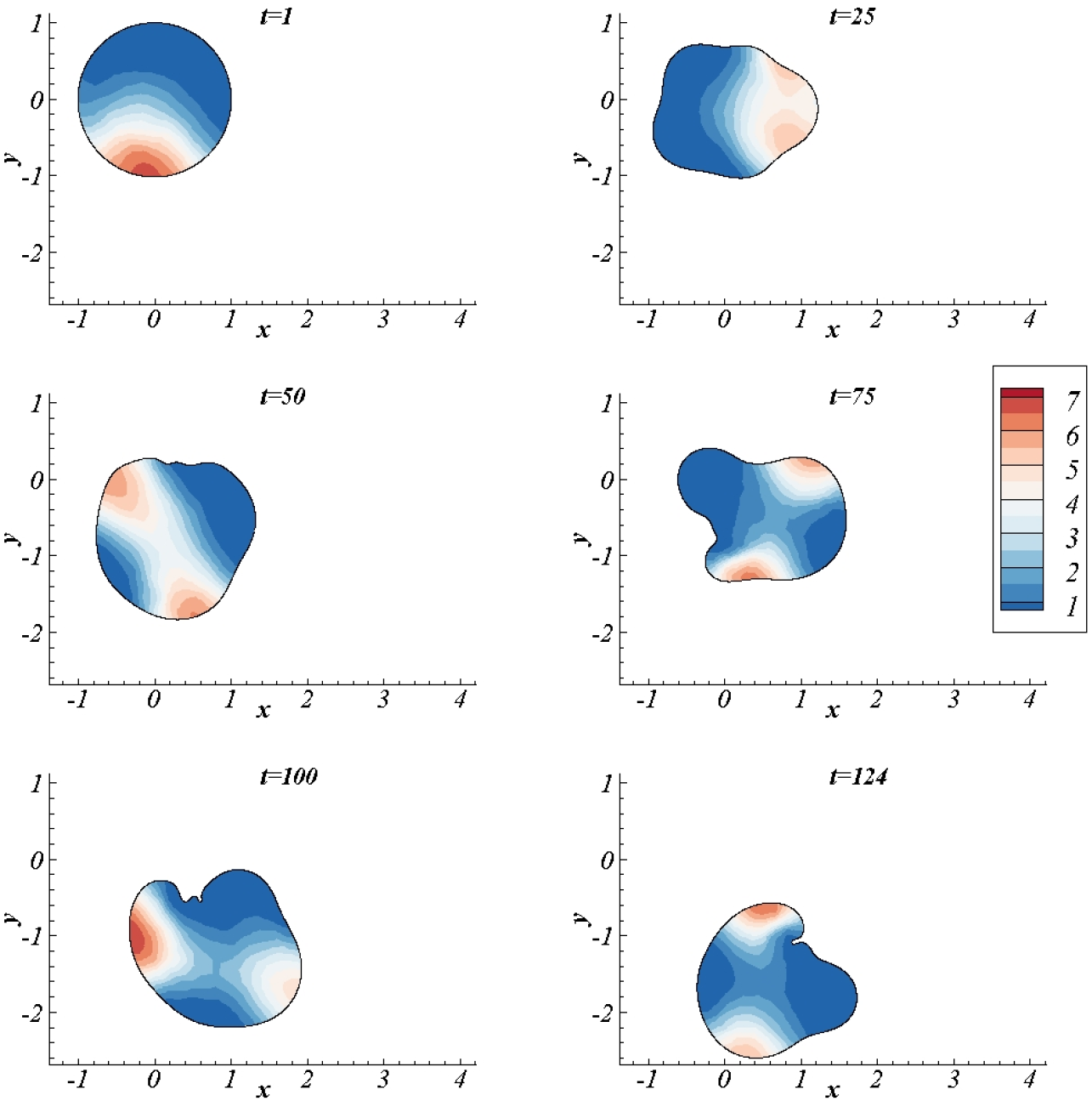
Results of spontaneous cell migration. At *t* = 1, 25, 50, 75,100,124. The colour map indicates the *a_b_* field.

Lastly, Section 6(a) describes the quality of the mesh throughout the simulation. Although the maximum quality was again 1, the average and the minimum qualities fluctuated more and reached lower values compared to the chemotactic case. Furthermore, Section 6(b) shows two stages. At the beginning, until about *t* = 20, the smoothing scheme was enough to 730 deal with the mesh deformation. However, after *t* = 20, it was not enough 731 and many more runs of the re-meshing scheme were needed.

**Fig 8.**
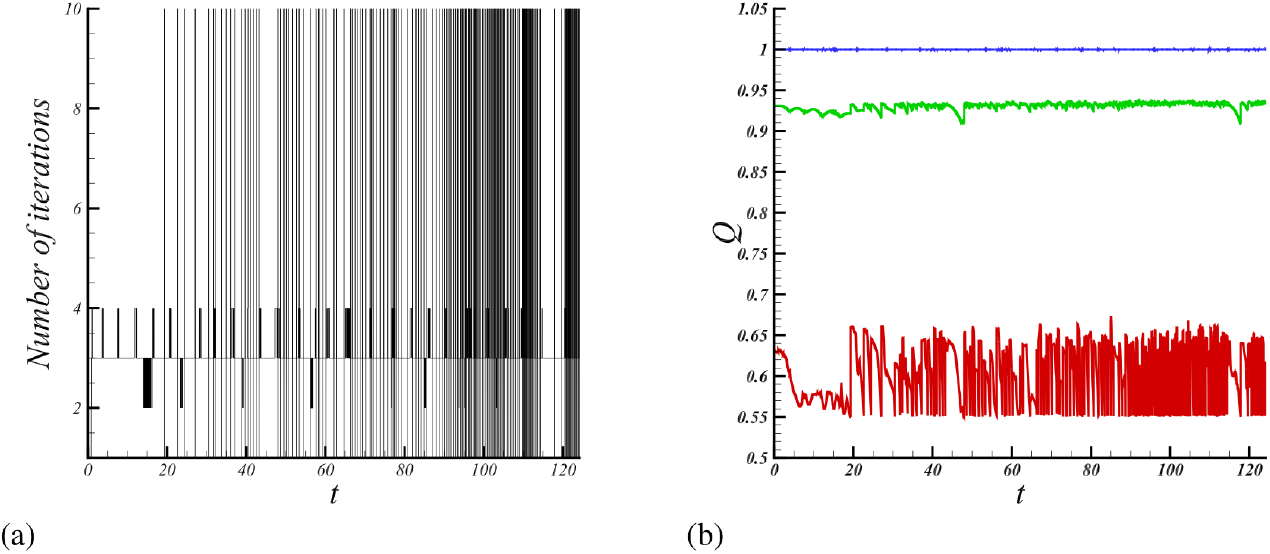
Report of mesh statistics and smoothing iterations at each step in spontaneous migration. (a) Statistics on the mesh regularity. From top to bottom the curves indicate respectively the maximum, the average and the minimum qualities. (b) Number of smoothing iterations. The vertical lines indicate a re-meshing step.

## 5 Discussion

In this study, we have formulated a mechanobiochemical model for single cell migration during directed and random migration. The model couples mechanical properties describing the mechanical structure of the cytoskeleton with biochemical processes describing the spatiotemporal dynamics of the actomyosin molecules during the process of single cell migration. We then employed a novel bulk-surface moving-mesh finite element method to discretise the model in space. Standard finite differences in time allowed us to solve the resulting semi-discrete weak variational forms. By employing in a novel way, smoothing and re-meshing algorithms, we are able to exhibit computationally large cell deformations during migration. The computational framework allows us to access easily geometric, mechanical and biochemical properties which play a crucial role during the process of single cell migration. Not only are we able to exhibit complex cell migration pathways as observed experimentally, we are also able to compute and quantify geometric quantities such as the curvature, material and mesh velocities, area, speed, cell stiffness and the spatiotemporal distribution of the actomyosin molecules. This study significantly contributes to an emerging area of investigating the role of mechanics and biomolecular spatiotemporal dynamics during the process of single cell migration.

We briefly summarise here the computational modelling approach and its implementation. First, we saw that the Meinhardt model exhibits travelling–wave, chemotactic, and (ii) competing–peak, spontaneous random behaviours, along the cell membrane. Although the dynamics of the leading edge varied from cell to cell, most cells have travelling actin waves [31, 77]. These waves can appear locally or globally, depending on the cell and the extracellular environment on which it is migrating. For the case of chemotactically–driven migrating cells, the waves appear locally and help cells to avoid obstacles or reorientation or re-polarisation [31, 77]. Section 6 illustrates this phenomenon in human breast cancer (MDA–MB–231 line). By following the arrows, which indicate the position of the leading edge, we appreciate the travelling wave from the top left to the bottom right of Section 6. This phenomenon, where the leading edge travels from one location to another, is successfully mimicked in the mechanobiochemical model in the presence of chemotractant as illustrated in Section 6(a).

**Fig 9.**
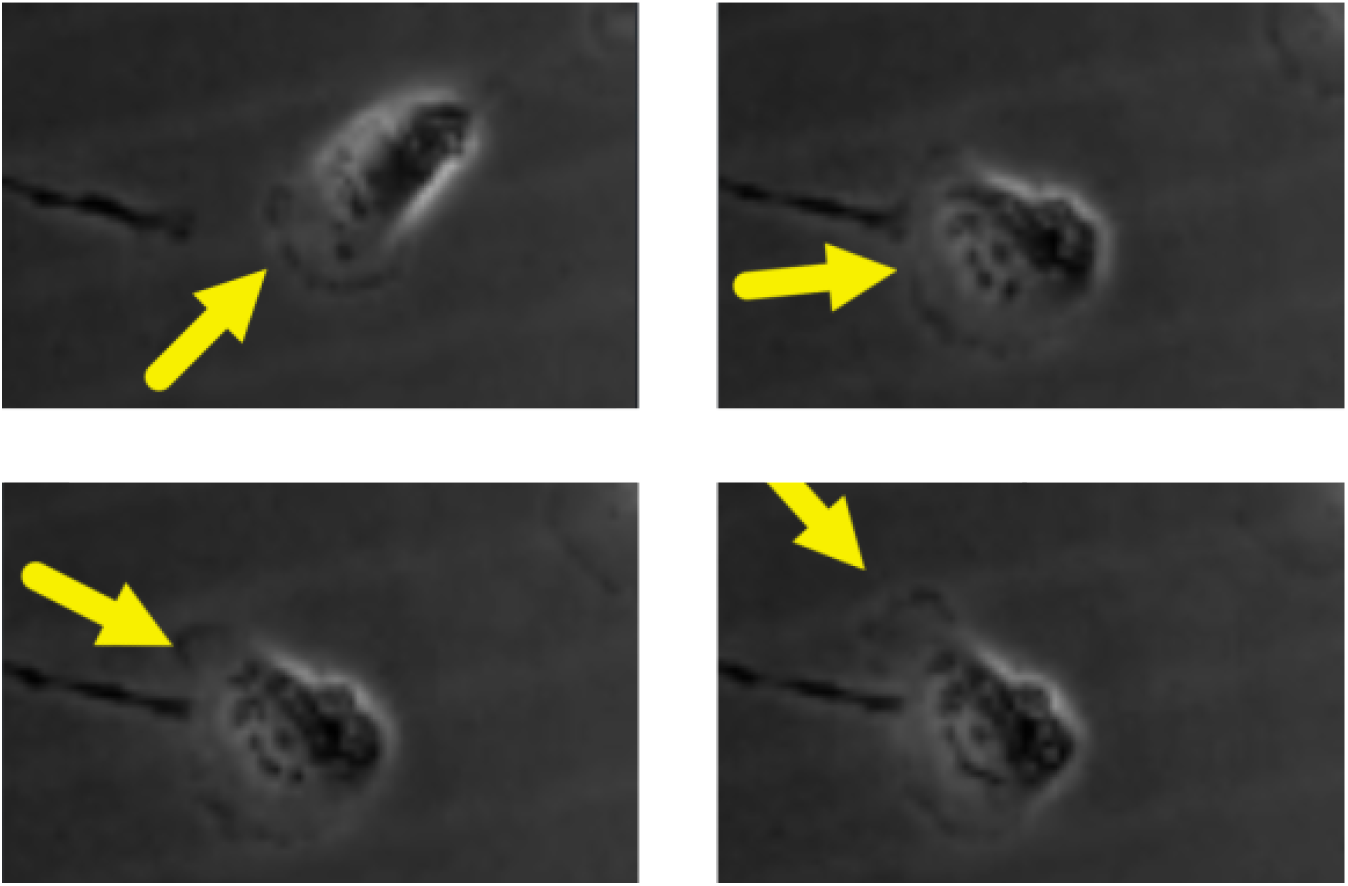
Picture shots from a cell migration assay. Experiment using human breast cancer (MDA-MB-231 line) presented in [72] (permission given by Dr. Gau). The arrows point to the leading edge and show how it travels along the membrane.

On the other hand, when cells move unsteadily the actin dynamics can include global travelling waves and the appearance of leading edges at different locations [31, 77]. In Section 6, the spontaneous–migration case reproduces this type of kinetic dynamics, as also reported in [26–28]. Although the response is similar, in the case presented here, the parameters are all homogeneous and thus the kinetics is completely spontaneous—in [26–28] the authors included a space–dependent chemotactic–signalling term. The dissimilarity in our chemotactic response therefore is due to the difference in parameters and the signalling term. Nonetheless, even though the model can reproduce chemotactic and spontaneous migration, it utilises different mechanisms depending on the different parameters in each scenario; thus, a further analysis of the model is required. It would be very interesting to find a set of parameters able to provide a local travelling wave in chemotactic migration as well as competing peaks in spontaneous migration.

In addition, our results show a marked round–like shape. This is common in amoeboid movement which is characterised by small or highly homogeneously distributed adhesion points and strong actomyosin–mediated contractility [73]. In our mechanobiochemical model, the adhesion points are represented by the right–hand–side term of Eq. (10) which is space–independent. The contraction, dominated by *λ*(*t*), is also homogeneous and therefore does not precisely model actomyosin–mediated contractility because *λ* is not polarised. Yet, it does contribute to maintain the round shape of the domain. Additionally, at time *t* = 94 for the chemotactic case, we see that the rear has a small protrusion (Section 6(a)) that can be also appreciated in Section 6. Although the protrusion in the simulation is not as large as the one observed in experiments [72], it indicates qualitative agreements with experiments.

In the case of spontaneous migration, the trajectory (Section 6) exhibits zigzag turning and some directional persistence. Da Yang et al. [74] describes this phenomenon for microglia cells from rats (PMG) and mice (MG5). In their experiments, cells were able to move freely, without external guidance and at a very low density, showing a tendency to advance in an almost linear straight direction followed by a sudden change in direction. Moreover, they determined a statistically significant zigzag pattern. In short, they showed that it is more likely that a microglia cell turn in the opposite direction of the previous turn than in the same direction. In other words, after turning with a positive angle it is more likely that the cell will turn with a negative angle and vice versa. In addition, they also reported a short–term directional persistence. This response of the Meinhardt model has been previously reported in [26, 27].

## 6 Conclusions

In this work a computational modelling framework for the simulation of cell migration was introduced. The proposed framework is able to mimic both spontaneous and chemotactic or directed single cell migration with an appropriate selection of parameters. In addition, this computational framework deals with highly deforming domains by means of three algorithms: the De Boor’s [67] algorithm to equidistribute the boundary mesh, the Durand’s [68] algorithm to smooth the bulk mesh, and the Engwirda’s [69, 70] toolbox to remesh the domain when necessary. Furthermore, it is able to model and simulate bulk–surface processes that drive single cell migration on evolving two–dimensional domains.

In the current work, we presented a bulk–surface moving–mesh finite element method that allows the computation of a deforming domain for long periods of time and a simplified model for cell migration. In addition, we also improved the Durand’s [68] algorithm by combining it with the De Boors’s [67] algorithm. Durand et al. [68] indicated that non–straight surface meshes, for example a circle, could not be smoothed with their algorithm; however, the De Boor’s [67] algorithm presented here, easily deals with any sort of surface.

In short, our model considers a simplified biological system where the information goes from the membrane kinetics to the cytosolic kinetics that then leads to a mechanical response. Furthermore, as the mechanical response deforms the geometry—which at the same time affects the dynamics on the membrane and in the bulk—it indirectly affects the biochemical system. The membrane–protein dynamics is modelled with the Meinhardt’s [33] system for cell orientation, which can evolve in spontaneous competing peaks or in a forced and directed peak. This last response appears as an amplification of a space–dependent reaction rate that we compare to the activity of membrane receptors. The downstream effect from the plasma membrane to the cytoskeleton is given by a diffusion-depletion system that assumes that the actin–filament density close to the membrane is proportional to activity of the membrane proteins. Thus, we considered that while the membrane activity affects the cytosolic activity, the latter does not affect the former. Finally, the model includes a mechanical response from the cytoskeleton activity. Mathematically, the mechanical model can be regarded as the product of: a local isotropic expansion due to the growth of actin filaments, a global contraction due to the area change, a homogeneous attachment to the substratum and curvature–dependent forces on the contour. Under the biological perspective, expansion and contraction occur as a consequence of the dynamics of the actin cytoskeleton and myosin proteins. In addition, elastic supports refer to focal adhesions and the curvature–dependent forces represent the membrane tension.

This study is a first step towards the formulation of a comprehensive predictive mechanobiochemical model for single cell migration where we couple mechanical and biochemical processes using the the bulk (cytosol) and surface (cortex) spatiotemporal dynamics approach. Several improvements to the model are necessary is it is to be useful for experimental purposes. For example, the use of dummy parameters, the type of material behaviour, the lack of direct interpretation of the kinetic species, the required computational time, among others, need further studies.

Our modelling philosopher is that due to the complexity of the biological problem of single cell migration, the best approach to build a robust cell migration simulator in a modular way. In other words, developing each stage of the information flow separately and consistently with the biological theory and after that combining the modules with proper cross-communication. Thus, future work could be focused on different modular aspects such as:

- more complex mechanical models, for example, the implementation of visco-elasticity or hyper-elasticity theories, to better describe the dynamics of the cytoskeleton,
- the development of biologically-inspired cytosolic and membrane kinetics and communication, for example, the formulation of bulk-surface PDEs where the variables directly represent specific molecules,
- different ways to model focal adhesions, for instance, since they do not appear everywhere and are constantly degraded, a stochastic equation could describe their location and a reaction equation for their strength, **Ψ**,
- a more discrete modelling of the cytosolic and membrane kinetics, for example, active and inactive GPCR’s could be regarded as discrete entities able to modify the actin cytoskeleton at specific locations,
- the inclusion of the bending forces on the membrane,
- the use of experimental data to fit the model parameters, and
- introducing the extracellular matrix to study how single cells migrate through complex non-isotropic environments.

## Acknowledgments

DHA was supported by the Universidad Nacional de Colombia through resolutions: 405 of 2019, 051 and 0354 of 2020, and 166 of 2021. DHA thanks the University of Sussex for its hospitality during his one–month research visit to the UK. AM is partly supported by the EPSR grant number EP/J016780/1, the European Union Horizon 2020 research and innovation programme under the Marie Sklodowska-Curie grant agreement No 642866, the Commission for Developing Countries, and the Simons Foundation. AM acknowledges the Royal Society Wolfson Research Merit Award (2016-2021) funded generously by the Wolfson Foundation.

## References

1. Camley BA, Zhao Y, Li B, Levine H, Rappel WJ. Crawling and turning in a minimal reaction-diffusion cell motility model: Coupling cell shape and biochemistry. Physical Review E. 2017;95(1):1–13. doi:10.1103/PhysRevE.95.012401.

2. Seetharaman S, Etienne-Manneville S. Cytoskeletal Crosstalk in Cell Migration. Trends in Cell Biology. 2020;30(9):720–735. doi:10.1016/j.tcb.2020.06.004.

3. Hobson CM, Stephens AD. Modeling of Cell Nuclear Mechanics: Classes, Components, and Applications. Cells. 2020;9(7). doi:10.3390/cells9071623.

4. Yamada KM, Sixt M. Mechanisms of 3D cell migration. Nat Rev Mol Cell Biol. 2019;20:738–752. doi:10.1038/s41580-019-0172-9.

5. Alberts B, Johnson A, Lewis J, Morgan D, Raff M, Roberts K, et al. The Cytoskeleton. In: Molecular Biology of the Cell. 6th ed. Garland Science; 2015. p. 880–962.

6. Uriu K, Morelli LG, Oates AC. Interplay between intercellular signaling and cell movement in development. Seminars in Cell and Developmental Biology. 2014;35:66–72. doi:10.1016/j.semcdb.2014.05.011.

7. Morales T. Chondrocyte Moves: clever strategies ? Osteoarthritis Cartilage. 2007;15(8):861–871.

8. Othmer HG. Eukaryotic cell dynamics from crawlers to swimmers. Wiley Interdisciplinary Reviews: Computational Molecular Science. 2019;9(1):e1376. doi:10.1002/wcms.1376.

9. Warner H, Wilson BJ, Caswell PT. Control of adhesion and protrusion in cell migration by Rho GTPases. Current Opinion in Cell Biology. 2019;56:64–70. doi:10.1016/j.ceb.2018.09.003.

10. Shah EA, Keren K. Mechanical forces and feedbacks in cell motility. Current Opinion in Cell Biology. 2013;25(5):550–557. doi:10.1016/j.ceb.2013.06.009.

11. Ridley AJ, Schwartz MA, Burridge K, Firtel RA, Ginsberg MH, Borisy G, et al. Cell Migration: Integrating Signals from Front to Back. Science. 2003;302(5651):1704–1709. doi:10.1126/science.1092053.

12. Buttenschön A, Edelstein-Keshet L Bridging from single to collective cell migration: A review of models and links to experiments. PLOS Computational Biology. 2020;16(12): e1008411. doi:10.1371/journal.pcbi.1008411.

13. Murray JD. Mathematical Biology: I. An Introduction. 3rd ed. Springer; 2002.

14. Murray JD. Mathematical Biology II: Spatial Models and Biomedical Applications. 3rd ed. Springer; 2003.

15. Harris PJ. A simple mathematical model of cell clustering by chemotaxis. Mathematical Biosciences. 2017;294(May):62–70. doi:10.1016/j.mbs.2017.10.008.

16. Holmes WR, Park J, Levchenko A, Edelstein-Keshet L A mathematical model coupling polarity signaling to cell adhesion explains diverse cell migration patterns. PLOS Computational Biology. 2017;13(5):e1005524. doi:10.1371/journal.pcbi.1005524.

17. Gonçalves IG, Garcia-Aznar JM Extracellular matrix density regulates the formation of tumour spheroids through cell migration. PLOS Computational Biology. 2021;17(2):e1008764. doi:10.1371/journal.pcbi.1008764.

18. Zhao J, Cao Y, Dipietro LA, Liang J. Dynamic cellular finite-element method for modelling large-scale cell migration and proliferation under the control of mechanical and biochemical cues: a study of re-epithelialization. J R Soc Interface. 2017;14(129). doi:10.1098/rsif.2016.0959.

19. Alt W, Tranquillo RT. Basic morphogenetic system modeling shape changes of migrating cells, how to explain fluctuating lamellipodial dynamics. Journal of Biological Systems. 1995;3(4):905–916.

20. Séguis JC, Burrage K, Erban R, Kay D. Simulation of cell movement through evolving environment: a fictitious domain approach. University of Oxford; 2012.

21. Stéphanou A, Tracqui P. Cytomechanics of cell deformations and migration: from models to experiments. C R Biologies. 2002;325:295–308.

22. Fuhrmann J, Käs J, Stevens A. Initiation of cytoskeletal asymmetry for cell polarization and movement. Journal of Theoretical Biology. 2007;249:278–288. doi:10.1016/j.jtbi.2007.08.013.

23. Mori Y, Jilkine A, Edelstein-Keshet L. Wave-pinning and cell polarity from a bistable reaction-diffusion system. Biophysical Journal. 2008;94(9):3684–3697. doi:10.1529/biophysj.107.120824.

24. Cusseddu D, Edelstein-Keshet L, Mackenzie JA, Portet S, Madzvamuse A. A coupled bulk-surface model for cell polarisation. Journal of Theoretical Biology. 2019;481:119–135. doi:10.1016/j.jtbi.2018.09.008.

25. Mackenzie, J., Rowlatt, C. and Insall, R. A conservative finite element ALE scheme for mass-conservative reaction-diffusion equations on evolving two-dimensional domains. SIAM Journal on Scientific Computing, 43(1), pp.B132–B166.

26. Neilson MP, Mackenzie JA, Webb SD, Insall RH. Modeling cell movement and chemotaxis using pseudopod-based feedback. Computational Methods in Science and Engineering. 2011;33(1):1035–1057.

27. Elliott CM, Stinner B, Venkataraman C. Modelling cell motility and chemotaxis with evolving surface finite elements. J R Soc Interface. 2012;9(June):3027–3044.

28. Campbell EJ, Bagchi P, Campbell EJ, Bagchi P. A computational model of amoeboid cell swimming A computational model of amoeboid cell swimming. Physics of Fluids. 2017;29:101902:1–101902:16.

29. Cheng Y, Othmer H. A Model for Direction Sensing in Dictyostelium discoideum: Ras Activity and Symmetry Breaking Driven by a Gβγ-Mediated, Gα2-Ric8 – Dependent Signal Transduction Network. PLoS Computational Biology. 2016;12(5):e1004900. doi:10.1371/journal.pcbi.1004900.

30. Bhattacharya S, Iglesias PA. The Regulation of Cell Motility Through an Excitable Network. IFAC PapersOnLine. 2016;49(26):357–363. doi:10.1016/j.ifacol.2017.03.001.

31. Allard J, Mogilner A. Traveling waves in actin dynamics and cell motility. Current Opinion in Cell Biology. 2013;25(1):107–115. doi:10.1016/j.ceb.2012.08.012.

32. Goehring NW, Grill SW. Cell polarity: mechanochemical patterning. Trends in Cell Biology. 2013;23(2):72–80. doi:10.1016/j.tcb.2012.10.009.

33. Meinhardt H. Orientation of chemotactic cells and growth cones: models and mechanisms. Journal of Cell Science. 1999;112:2867–2874.

34. SJ Han, S Kwon, K Sook Kim Contribution of mechanical homeostasis to epithelial–mesenchymal transition. Cellular Oncology. 2022;45:1119–1136. doi:10.1007/s13402-022-00720-6.

35. JA Espina, CL Marchant, EH Barriga Durotaxis: the mechanical control of directed cell migration. The FEBS Journal. 2022;289:2736–2754. doi:10.1111/febs.15862.

36. F Merino-–Casallo, MJ Gomez–Benito, S Hervas–Raluy, JM Garcia–-Aznar Durotaxis: the mechanical control of directed cell migration. The FEBS Journal. 2022;289:2736–2754. doi:10.1111/febs.15862.

37. Lewis MA, Murray JD. Analysis of stable two-dimensional patterns in contractile cytogel. J Nonlinear Sci. 1991;1:289–311. doi:10.1007/BF01238816.

38. Murphy L, Madzvamuse A. A moving grid finite element method applied to a mechanobiochemical model for 3D cell migration. Applied Numerical Mathematics. 2020;158:336–359. doi:10.1016/j.apnum.2020.08.004.

39. Madzvamuse A, George UZ. The moving grid finite element method applied to cell movement and deformation. Finite Elements in Analysis and Design. 2013;74:76–92. doi:10.1016/j.finel.2013.06.002.

40. Dziuk G, Elliott CM. Finite elements on evolving surfaces. IMA Journal of Numerical Analysis. 2007;27(2):262–292. doi:10.1093/imanum/dri017.

41. Vorotnikov AV. Chemotaxis: Movement, Direction, Control. Biochemistry (Moscow). 2011;76(13):1528–1555.

42. Artemenko Y, Lampert TJ, Devreotes PN. Moving towards a paradigm: common mechanisms of chemotactic signaling in Dictyostelium and mammalian leukocytes. Cellular and molecular life sciences: CMLS. 2014;71(19):3711–3747. doi:10.1007/s00018-014-1638-8.

43. Krause M, Gautreau A. Steering cell migration: lamellipodium dynamics and the regulation of directional persistence. Nature Reviews Molecular Cell Biology. 2014;15(9):577–590. doi:10.1038/nrm3861.

44. Devreotes P, Horwitz AR. Signaling Networks that Regulate Cell Migration. Cold Spring Harbor Perspectives in Biology. 2015;7(8):a005959. doi:10.1101/cshperspect.a005959.

45. Onsum MD, Rao CV. Calling heads from tails: the role of mathematical modeling in understanding cell polarization. Current Opinion in Cell Biology. 2009;21(1):74–81. doi:10.1016/j.ceb.2009.01.001.

46. Rappel WJ, Edelstein-Keshet L. Mechanisms of cell polarization. Current Opinion in Systems Biology. 2017;3:43–53. doi:10.1016/j.coisb.2017.03.005.

47. Irgens F. Theory of Elasticity. In: Continuum Mechanics. Berlin, Heidelberg: Springer Berlin Heidelberg; 2008. p. 199–302.

48. Ferreira AJM. Plane stress. In: MATLAB Codes for Finite Element Analysis. Solid Mechanics and its Applications. vol. 157 of Solid Mechanics and its Applications. Dordrecht: Springer, Dordrecht; 2009. p. 143–152.

49. Barreira R, Elliott CM, Madzvamuse A. Mathematical Biology The surface finite element method for pattern formation on evolving biological surfaces. J Math Biol. 2011;63:1095–1119. doi:10.1007/s00285-011-0401-0.

50. Frittelli M, Madzvamuse A, Sgura I, Venkataraman C. Numerical Preservation of Velocity Induced Invariant Regions for Reaction – Diffusion Systems on Evolving Surfaces. J Sci Comput. 2018;77(2):971–1000. doi:10.1007/s10915-018-0741-7.

51. Dziuk G, Elliott CM. Finite element methods for surface PDEs. Acta Numerica. 2013;22(April):289–396. doi:10.1017/S0962492913000056.

52. Wang Y, Irvine DJ. Convolution of chemoattractant secretion rate, source density, and receptor desensitization direct diverse migration patterns in leukocytes. Integrative Biology. 2013;5(3):481–494. doi:10.1039/c3ib20249f.

53. Madzvamuse A, Maini PK, Wathen AJ. A moving grid finite element method for the simulation of pattern generation by turing models on growing domains. Journal of Scientific Computing. 2005;24(2):247–262. doi:10.1007/s10915-004-4617-7.

54. Stéphanou A, Chaplain MAJ, Tracqui P. A mathematical model for the dynamics of large membrane deformations of isolated fibroblasts. Bulletin of Mathematical Biology. 2004;66(5):1119–1154. doi:10.1016/j.bulm.2003.11.004.

55. George UZ, Stéphanou A, Madzvamuse A. Mathematical modelling and numerical simulations of actin dynamics in the eukaryotic cell. Journal of Mathematical Biology. 2013;66(3):547–593. doi:10.1007/s00285-012-0521-1.

56. Heine CJ. Isoparametric finite element approximation of curvature on hypersurfaces. Preprint, University Freiburg. 2004;0:1–17. doi:10.1093/imanum/drs037.

57. Barrett JW, Garcke H, Nüurnberg R. Chapter 4 - Parametric finite element approximations of curvature-driven interface evolutions. In: Bonito A, Nochetto RHBTHoNA, editors. Geometric Partial Differential Equations - Part I. vol. 21. Elsevier; 2020. p. 275–423.

58. Novak IL, Gao F, Choi YS, Resasco D, Schaff JC, Slepchenko BM. Diffusion on a curved surface coupled to diffusion in the volume: Application to cell biology. Journal of Computational Physics. 2007;226(2):1271–1290. doi:10.1016/j.jcp.2007.05.025.

59. Rätz A, Röger M. Symmetry breaking in a bulk-surface re-action-diffusion model for signalling networks. Nonlinearity. 2014;27(8):1805–1827. doi:10.1088/0951-7715/27/8/1805.

60. Rätz A. Turing-type instabilities in bulk-surface reaction–diffusion systems. Journal of Computational and Applied Mathematics. 2015;289:142–152. doi:10.1016/j.cam.2015.02.050.

61. Elliott CM, Ranner T, Venkataraman C. Coupled Bulk-Surface Free Boundary Problems Arising from a Mathematical Model of Receptor-Ligand Dynamics. SIAM Journal on Mathematical Analysis. 2017;49(1):360–397. doi:10.1137/15M1050811.

62. Elliott CM, Ranner T. Finite element analysis for a coupled bulk-surface partial differential equation. IMA Journal of Numerical Analysis. 2013;33(2):377–402. doi:10.1093/imanum/drs022.

63. Alhazmi M. Exploring Mechanisms for Pattern Formation through Coupled Bulk-Surface PDEs in Case of Non-linear Reactions. International Journal of Advanced Computer Science and Applications. 2019;10(3):556–568. doi:10.14569/IJACSA.2019.0100372.

64. Frittelli M, Madzvamuse A, Sgura I. Bulk-surface virtual element method for systems of PDEs in two-space dimensions. Numerische Mathematik. 2021;147(2):305–348. doi:10.1007/s00211-020-01167-3.

65. Brezzi F, Falk RS, Donatella Marini L. Basic principles of mixed Virtual Element Methods. ESAIM: Mathematical Modelling and Numerical Analysis. 2014;48(4):1227–1240. doi:10.1051/m2an/2013138.

66. Elliott CM, Styles V. An ALE ESFEM for solving PDEs on evolvoing surfaces. Milan J Math. 2012;80:469–501.

67. de Boor C. Good approximation by splines with variable knot. In: Numerical Solution of Differential Equations. Dundee: Lecture Notes in Math. 363, Springer, 1974; 1973. p. 12–20.

68. Durand R, Pantoja-rosero BG, Oliveira V. A general mesh smoothing method for finite elements. Finite Elements in Analysis Design. 2019;158(February):17–30. doi:10.1016/j.finel.2019.01.010.

69. Engwirda D. Unstructured mesh methods for the Navier-Stokes equations; 2005.

70. Engwirda D. Locally-optimal Delaunay-refinement and optimisation-based mesh generation. The University of Sydney; 2014.

71. Campillo-Funollet E, Venkataraman C, Madzvamuse A. Bayesian Parameter Identification for Turing Systems on Stationary and Evolving Domains. Bulletin of Mathematical Biology. 2019;81(1):81–104. doi:10.1007/s11538-018-0518-z.

72. Gau D, Roy P. Single Cell Migration Assay Using Human Breast Cancer MDA-MB-231 Cell Line. Bio-protocol. 2020;10(8):e3586. doi:10.21769/BioProtoc.3586.

73. Friedl P, Alexander S. Cancer invasion and the microenvironment: Plasticity and reciprocity. Cell. 2011;147(5):992–1009. doi:10.1016/j.cell.2011.11.016.

74. Da Yang T, Park JS, Choi Y, Choi W, Ko TW, Lee KJ. Zigzag turning preference of freely crawling cells. PLoS ONE. 2011;6(6):e20255. doi:10.1371/journal.pone.0020255.

75. Cusseddu, D. and Madzvamuse, A. Numerical investigations of the bulk-surface wave pinning model Mathematical Biosciences, 2022, 354, 108925

76. Madzvamuse, A., and Chung, A. H. The bulk-surface finite element method for reaction–diffusion systems on stationary volumes Finite Elements in Analysis and Design, 2016, 108, 9–21.

77. Kamps, D., Koch, J., Juma, V.O., Campillo-Funollet, E., Graessl, M., Banerjee, S., Mazel, T., Chen, X., Wu, Y.W., Portet, S. and Madzvamuse, A. Optogenetic tuning reveals rho amplification-dependent dynamics of a cell contraction signal network. Cell reports, 2020, 33(9), p.108467.

